# NFYA promotes the malignant behavior of triple-negative breast cancer through the regulation of lipid metabolism

**DOI:** 10.1101/2022.05.26.493660

**Authors:** Nobuhiro Okada, Chihiro Ueki, Masahiro Shimazaki, Goki Tsujimoto, Susumu Kohno, Hayato Muranaka, Kiyotsugu Yoshikawa, Chiaki Takahashi

## Abstract

Two splicing variants exist in NFYA, which exhibits high expression in many human tumor types, and their expression balance is known to correlate with prognosis in breast cancer, but functional differences are still unclear. Here, we demonstrate that NFYAv1, the long-form variant, upregulates the transcription of ACACA and FASN, essential lipogenic enzymes, to enhance the malignant behavior of triple-negative breast cancer (TNBC). Loss of the NFYAv1-lipogenesis axis strongly suppresses the malignant behavior in vitro and in vivo, indicating that the NFYAv1-lipogenesis axis is essential for TNBC malignant behavior and that the axis might be a potential therapeutic target for TNBC. Furthermore, mice deficient in lipogenic enzymes such as Acly, Acaca, and Fasn exhibit embryonic lethality, but our Nfyav1-deficient mice have no apparent developmental abnormalities. Taken together, our results elucidate NFYAv1-lipogenesis axis has significant tumor-promoting effects and the potential for NFYAv1 to be a safer therapeutic target for TNBC.

## Introduction

Triple-negative breast cancer (TNBC) is an aggressive breast cancer subtype with a lack of the expression of estrogen receptor (ER), progesterone receptor (PgR), and HER2 protein^1^. TNBC is sensitive to chemotherapy but not to endocrine therapy and has no identified molecular targets, resulting in an overall poor prognosis. Therefore, there is an urgent need to develop new treatment regimens or identify molecular targets for therapy. Recent remarkable advances in cancer metabolism research have revealed that reprogramming lipid metabolism is one of the hallmarks of cancer^2, 3^. It has been reported that lipogenic enzymes are highly expressed in TNBC to obtain membrane phospholipids, signaling molecules, protein modifications, and energy for high proliferation^2, 4–7^. Therefore, preclinical and clinical trials targeting lipid metabolism as a therapeutic target are proceeding^3, 8, 9^. However, there are still unknown regarding the selection of patients who will benefit from lipid metabolism blockade, drug resistance, and other issues. Furthermore, the more serious problem is that knockout mice of the lipogenic enzymes, Acly, Acaca, or Fasn, have been reported to show embryonic lethality in early embryogenesis, raising safety concerns about considering the lipogenesis pathway blockade by inhibiting these genes as a therapeutic target^10–12^. Therefore, an important issue is to identify a safer target that can be expected to have therapeutic effects comparable to inhibition of those enzymes.

NF-Y is a trimeric protein complex composed of three subunits, NFYA, NFYB, and NFYC, that binds to the CCAAT box on the promoter of target genes to activate transcription^13, 14^. NFYA is considered a limiting subunit of the trimeric complex because it has a DNA-binding domain at the C-terminus, and point mutations in the DNA binding domain inhibit NF-Y-DNA complex formation^15^. There are two types of splice variants of NFYA: long-form (NFYAv1) and short-form (NFYAv2) that lacks the 29 amino acids encoded by exon 3 in the Q-rich transactivation domain^16^. The genome-wide studies have shown that CCAAT boxes and NF-Y binding are enriched at the promoters of genes overexpressed in various types of cancer, thereby eliciting various biological effects, including metabolic regulation^13, 17^. Clinical studies have shown that patients with overexpression of NF-Y target genes in several types of cancer have a poor prognosis^13, 18, 19^. Integrative analysis of NF-Y genome binding and gene expression profiles by knockdown of NF-Y subunits revealed that NF-Y regulates de novo biosynthetic pathways of lipids^17^. Such NF-Y effects on metabolic regulation in the lipogenesis are exerted in concert with sterol regulatory element-binding protein1 (SREBP1), and a genome-wide scan of SREBP1 and NF-Y occupancy showed that NF-Y occupies 32 % of SREBP1 targets in human hepatocyte cells^20^. NF-Y also activates glycolytic genes, represses mitochondrial respiratory genes, and targets genes involved in the SOCG (serine, one-carbon, glycine) pathway, glutamine pathway, and polyamines and purines biosynthesis, indicating that metabolic pathways are globally altered and specific cancer-causing nodes are under NF-Y control^17, 21^. A high-level NFYA expression has been detected in various human cancer types, including breast, prostate, gastric adenocarcinoma, and lung squamous cell carcinomas^22–25^. The expression balance of two splice variants of NFYA varies in a tissue- and cell-specific manner, and a correlation between the expression balance and prognosis has been reported in breast cancer^16, 23^. Thus, it was reported that the involvement of NF-Y in oncogenic mechanism and metabolic pathway regulation; however, these studies are based primarily on systematic investigations of global gene expression in large cohorts of cancer patients and cancer cells. The detailed function of NFYA in cancer carcinogenesis and malignant progression based on genetic and biochemical methods remains unclear.

This study investigated the functional importance of the NFYA splicing variants in TNBC. Using in vitro and in vivo models, we demonstrated that NFYAv1 promotes the malignant behavior of TNBC by enhancing transcription of the lipogenic enzymes, ACACA and FASN. Furthermore, our Nfyav1-specific knockout mice in this study show tumor suppression via suppression of ACACA and FASN in breast cancer tissues without any apparent developmental abnormalities. Altogether, our studies provide insight into the tumor-promoting effects of the NFYAv1-lipogenesis axis and indicate that NFYAv1 may be helpful as a therapeutic marker and target for TNBC.

## Results

### NFYA switches the expression of alternative splicing variants during EMT progression

Emerging evidence suggests the splicing variants of NFYA as essential components of the oncogenic signals in the development of breast cancers^23, 26^. In this study, we also found that breast cancer cells isolated from mouse breast tumor tissue significantly increased the expression of both variants of NFYA compared with non-transformed mouse mammary epithelial cells (NMuMG cells) (Fig. 1b, c). Although these results suggest that it is promising that NFYA plays an essential function in the malignant behavior of breast cancer, the functional differences between NFYA variants remain unclear due to the absence of functional domains on the spliced exon (Fig. 1a). To investigate the functional differences between variants, we first examined the expression pattern of variants in various subtypes of breast cancer cells.

**Fig. 1.**
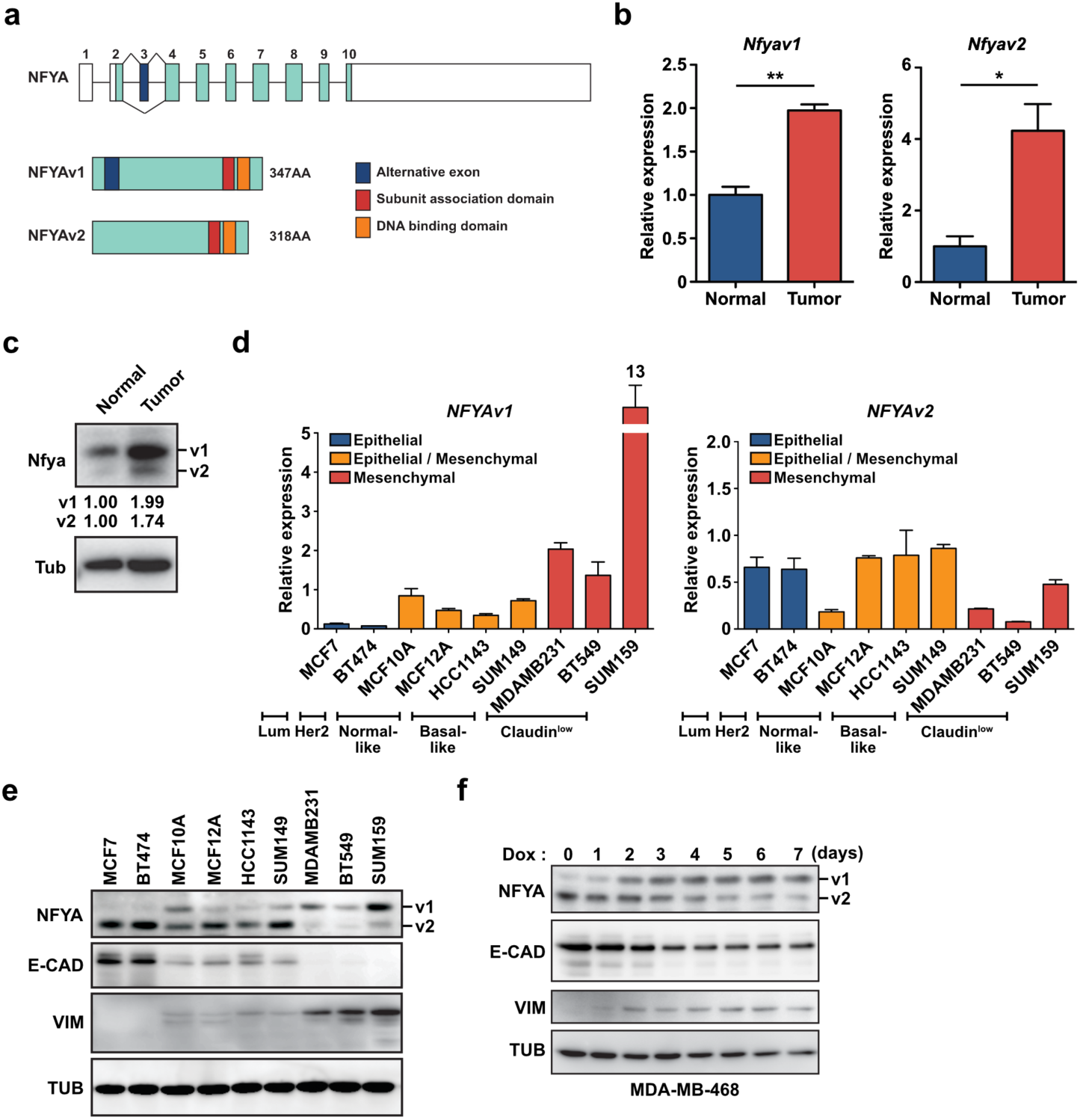
NFYA switches the expression of alternative splicing variants during EMT progression. **a** Schematic diagram of the intron-exon structure of *NFYA* gene. Alternative splicing of the third exon generates two different variants, long-form (v1) and short-form (v2). **b, c** qRT-PCR (**b**) and western blot analysis (**c**) of Nfyav1 and Nfyav2 in breast cancer cells (primary cells isolated from mouse breast cancer tissue) compared with non-transformed mouse mammary epithelial cells (NMuMG cells). **d** qRT-PCR analysis of *NFYAv1* (left panel) and *NFYAv2* (right panel) mRNA levels in various breast cancer cell lines. **e** Western blot analysis of NFYA, epithelial marker (E-CAD), and mesenchymal marker (VIM) protein levels in various breast cancer cell lines. **f** MDA-MB-468 cells were infected with lentivirus to express tet-on SNAIL and induced SNAIL expression by 1 µg/ml of doxycycline (Dox). Western blot analysis of NFYA protein level in the cells from day 0 to day 7 post-induction. E-CAD and VIM are markers for epithelial and mesenchymal cells, respectively. All error bars represent SEM; (*) P<0.05; (**) P<0.01.

Luminal and HER2-positive breast cancer cells predominantly express NFYAv2 but have almost undetectable NFYAv1, which is almost the same in normal-like and basal-like breast cancer cells NFYAv1 expression is a little detectable. On the other hand, Claudin^low^ breast cancer cells predominantly express NFYAv1 but have almost undetectable NFYAv2 (Fig. 1d, e). Interestingly, the expression pattern of NFYA is associated with epithelial-mesenchymal transition (EMT) status, in which NFYAv2 showed the same expression pattern as an epithelial marker, E-cadherin (CDH1), NFYAv1 showed the same as a mesenchymal marker, vimentin (VIM) (Fig. 1e; Supplementary Fig. 1a). An experiment using immortalized human mammary epithelial cells (HMLE cells) also proved the association between the expression of NFYA and EMT markers. We sorted CD44^low^/CD24^high^ (HMLE-Epi cells) and CD44^high^/CD24^low^ (HMLE-Mes cells) cells from HMLE cells and examined the EMT status and NFYA expression using sorted cells. Similar to the results of breast cancer cell lines, NFYAv1 expression correlates with vimentin contrasted to NFYAv2 correlates with E-cadherin (Supplementary Fig. 1b, c). Moreover, NFYA expression shifted from NFYAv2 to NFYAv1 in both MDA-MB-468 and HMLE cells manipulated EMT induction by overexpression of SNAIL (Fig. 1f; Supplementary Fig. 1d, e). Next, to examine whether the expression shift of NFYA is a reversible reaction, we treated SUM159 cells with Forskolin (FSK), known to induce mesenchymal-epithelial transition (MET)^27^. We found that the ratio of NFYAv2 to NFYAv1 increased through MET progression (Supplementary Fig. 1f). These results indicate that the expression of NFYA variants switches reversibly from NFYAv2 to NFYAv1 depending on EMT status, but NFYA itself is not an inducer of EMT because NFYA deficiency in HMLE cells had no effects on EMT progression (Supplementary Fig. 1g).

### NFYAv1 deficiency significantly inhibits tumor cell growth and tumorigenesis in TNBC

TNBC accounts for 10 to 15 % of all breast cancers, and TNBC therapy is still limited to chemotherapy because TNBC lacks ER, PgR, and HER2 expression, thereby having no sensitivity to endocrine therapy and anti-HER2 treatment. Therefore, TNBC has a worse prognosis than other breast cancers^1, 28, 29^. Therefore, to evaluate the potential of the NFYA-related pathway as a novel therapeutic target for TNBC, we focused mainly on the functional analysis of NFYA in TNBC, namely NFYAv1, to determine how it differs from NFYAv2. Given the increased expression of NFYA in breast cancer cells (Fig. 1b, c), we first generated NFYA-deficient TNBC cells by CRISPR/Cas9 system using SUM159 cells (Supplementary Fig. 2a-d) and examined whether NFYA contributes to the malignant behavior of TNBC. NFYA-deficient SUM159 cells showed a repressed carcinogenic phenotype, represented by a suppressed ability to grow the cells and form spheres and tumors (Fig. 2a-c). Re-expression of NFYAv1 in NFYA-deficient SUM159 cells significantly restored the malignant behavior. However, re-expression of NFYAv2 in NFYA-deficient SUM159 cells resulted in slight or no restoration of malignant behavior (Fig. 2d-f; Supplementary Fig. 2e). To further investigate NFYA variants’ requirements for malignant behavior of TNBC, we also generated NFYAv1-specific deficient SUM159 cells or overexpressed dominant-negative NFYA mutants in SUM159 cells. Though NFYAv1-specific deficient cells higher expressed NFYAv2 compared with control cells (Supplementary Fig. 2f), the cells suppressed the cell growth the same as NFYA-deficient cells (Fig. 2g). Dominant-negative mutants of NFYA have introduced three-point mutations into the DNA binding domain and repressed the ability of NF-Y as a transcription factor by losing the formation of the NF-Y-DNA complex^15^.

**Fig. 2.**
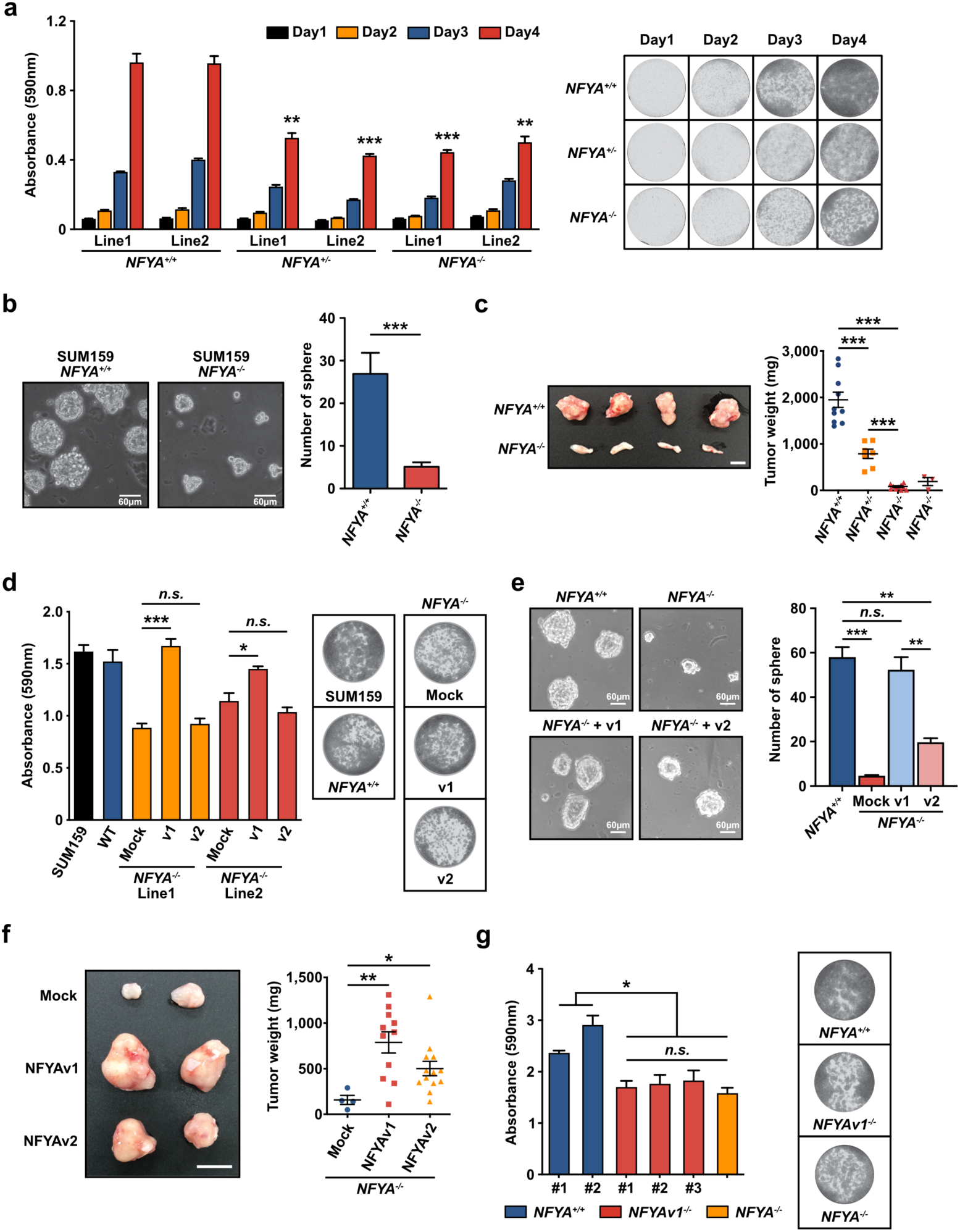
NFYAv1 deficiency significantly inhibits tumor cell growth and tumorigenesis in TNBC. **a** Quantification and representative pictures of 0.5 % crystal violet staining of *NFYA^+/+^*, *NFYA^+/-^*, and *NFYA^-/-^* SUM159 cells between days 1 and 4. **b** Representative images of sphere formation by *NFYA^+/+^*and *NFYA^-/-^* SUM159 cells. A bar graph shows the number of spheres larger than 60 µm in each group. **c** Representative images of tumors formed in mammary fat pads of NOD/SCID mice 50 days after implanted with *NFYA^+/+^* and *NFYA^-/-^*SUM159 cells. Dot plots show the weight of tumors from each experimental group (n=10; *NFYA^+/+^* and *NFYA^-/-^* (1st line), n=7; *NFYA^+/-^*, n=3; *NFYA^-/-^*(2nd line)). **d** Representative images of 0.5 % crystal violet staining of *NFYA^-/-^* SUM159 cells overexpressed each variant of NFYA on day 4. A bar graph shows the quantification of the staining. **e** Representative images of sphere formation by *NFYA^-/-^* SUM159 cells overexpressed each variant of NFYA. A bar graph shows the number of spheres larger than 60 µm in each group. **f** Representative images of tumors formed in mammary fat pads of NOD/SCID mice 50 days after implanted with *NFYA^-/-^* SUM159 cells overexpressed each variant of NFYA. Dot plots show the weight of tumors from each experimental group (n=4; Mock, n=11; NFYAv1, n=13; NFYAv2). **g** Representative images of 0.5 % crystal violet staining of two lines of *NFYA^+/+^*, three *NFYAv1^-/-^*, and *NFYA^-/-^* SUM159 cells on day 4. A bar graph shows the quantification of the staining. All error bars represent SEM; (*n.s.*) not significant; (*) P<0.05; (**) P<0.01; (***) P<0.001.

Overexpression of the dominant-negative mutant of NFYAv1 significantly suppressed the cell growth, whereas the effect of the dominant-negative mutant of NFYAv2 was only partial (Supplementary Fig. 2g). These observations suggest that NFYAv1, but not NFYAv2, is functionally important in the malignant behavior of TNBC. Furthermore, these effects might not be limited to TNBC but might affect other breast cancer subtypes as well because NFYA deficiency does not affect the cell growth and the ability of sphere formation in HMLE cells and luminal breast cancer cells, MCF7 cells, predominantly expressing NFYAv2 (Supplementary Fig. 1g, 2h-j).

### NFYA regulates lipid metabolism for the malignant behavior of TNBC

Some studies that analyzed global gene expression in cancer cell lines containing colon, liver, and lung cancer suggested that the NF-Y complex controls de novo lipogenesis with SREBPs^17, 20^. Furthermore, we also show that the expression of lipogenic enzymes, Acly, Acaca, and Fasn, increased in breast cancer cells, and inhibition of lipogenesis by FASN inhibitor, Cerulenin, completely inhibits TNBC cell growth and the sphere-forming ability (Supplementary Fig. 3a-c). These findings led us to hypothesize that NFYA could play an essential role in the malignant behavior of TNBC through the regulation of lipogenesis. To investigate the hypothesis, we first examined whether the repression of malignant behavior induced by NFYA deficiency could be attributed to lipid deficiency. The lipid mixture in the culture medium restored cell growth and the sphere-forming ability of NFYA-deficient SUM159 cells (Fig. 3a, b). Moreover, we found that NFYA deficiency dramatically reduced the number of lipid droplets, and the lipid mixture restored the number of lipid droplets (Fig. 3c). These results suggest that NFYA positively regulates lipogenesis for the malignant behavior of TNBC, and the loss of NFYA disrupts the mechanism. These phenotypic properties were also demonstrated by examining OCR (oxygen consumption rate) (Fig. 3d, e). NFYA wild-type SUM159 cells in normal conditions increased basal OCR by adding palmitic acid. In contrast, NFYA-deficient SUM159 cells did not differ the basal OCR level upon the addition of palmitic acid. Furthermore, in maximal respiratory conditions, maximal OCR in NFYA-deficient SUM159 cells treated with palmitic acid showed a very low OCR compared with NFYA wild-type SUM159 cells.

**Fig. 3.**
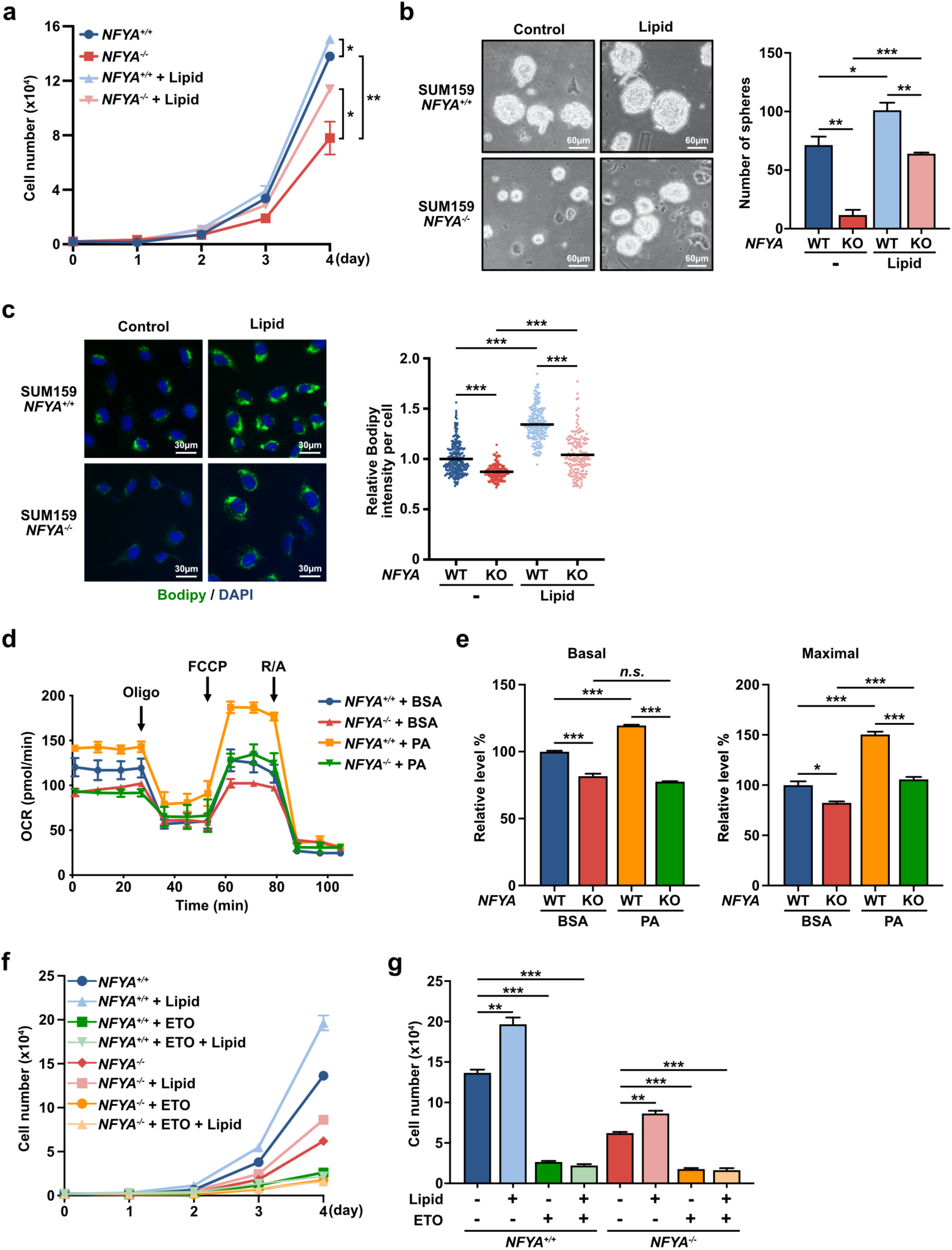
NFYA regulates lipid metabolism for malignant behavior of TNBC. **a** Cumulative population of cells was measured for 4 consecutive days in *NFYA^+/+^* and *NFYA^-/-^*SUM159 cells with or without the addition of lipid mixture. **B** Representative images of sphere formation by *NFYA^+/+^* and *NFYA^-/-^*SUM159 cells with or without the addition of lipid mixture. A bar graph shows the number of spheres larger than 60 µm in each group. **c** Representative fluorescence images of lipid droplet (green) detected with Bodipy 493/503 and nucleus (blue) detected with DAPI in *NFYA^+/+^*and *NFYA^-/-^* SUM159 cells with or without the addition of lipid mixture for 2 days. The dot plot shows the bodipy intensity per cell. Means represent SEM. (n=247; *NFYA^+/+^*, n=157; *NFYA^-/-^*, n=192; *NFYA^+/+^* with lipid, n=166; *NFYA^-/-^* with lipid). **d** Oxygen consumption rate (OCR) (mean ± SEM, n=3) in *NFYA^+/+^*and *NFYA^-/-^* SUM159 cells treated with the long-chain fatty acid palmitate (PA) or BSA. Oligo; oligomycin, FCCP; carbonyl cyanide-4-trifluoro methoxy phenyl hydrazone, R/A; Rotenone/antimycin A. **e** Experiments shown in Fig. 3d were quantified, and the relative levels of OCR associated with basal FAO and maximal FAO were calculated. **f** Cumulative population of cells was measured for 4 consecutive days in *NFYA^+/+^* and *NFYA^-/-^* SUM159 cells treated with or without lipid mixture, Etomoxir (ETO), and both. **g** A bar graph shows the cell number on day 4 shown in Fig. 3f. All error bars represent SEM; (*n.s.*) not significant; (*) P<0.05; (**) P<0.01; (***) P<0.001.

These results suggest that Fatty acid oxidation (FAO) does not occur in NFYA-deficient cells and is kept at a superficial level even under maximal respiratory conditions. However, we have not yet eliminated the possibility that NFYA deficiency disrupts glucose metabolism or the TCA cycle because acetyl-CoA, the source of fatty acids, is provided from those processes (Supplementary Fig. 3d)^30, 31^. Therefore, to eliminate the possibility, we evaluated the cell behavior by adding exogenous sodium acetate to supply acetyl-CoA without mitochondrial metabolism. As a result, the addition of sodium acetate had absolutely no effects on cell growth and accumulation of lipid droplets (Supplementary Fig. 3e, f). In addition, we found that NFYA-deficient cells were more sensitive to glucose deprivation than wild-type cells, which implies reduced lipid metabolism (Supplementary Fig. 3g). Finally, we evaluated cell growth with treatment by Etomoxir, an inhibitor of CPT1A, a critical rate-limiting enzyme of FAO, to corroborate that the restoration of malignant behavior in NFYA-deficient cells by adding lipid is indeed a response mediated by FAO. The results showed that inhibition of CPT1A completely abolished the effects of lipid addition (Fig. 3f, g).

### NFYA enhances lipogenesis by transcriptional activation of ACACA and FASN

Given that NFYA promotes the malignant behavior of TNBC by accelerating lipid synthesis, we next examined the expression of lipid metabolism-related genes to determine how NFYA regulates lipid metabolism, consisting of de novo lipogenesis and FAO involving multiple enzymatic reactions (Fig. 4a). In NFYA-deficient SUM159 cells, the expression of lipogenesis-related genes, including ACLY, ACACA, ACACB, and FASN, and FAO-related genes, including CPT1A and ACADL, were significantly decreased (Fig. 4b, c). Furthermore, the addition of lipids increased the expression of CPT1A and ACADL even in NFYA-deficient cells (Fig. 4b, c). Since the expression of ACLY and ACACB also increased by adding lipid mixture even in NFYA-deficient cells, we excluded those from the NFYA target and confirmed the protein level of ACACA and FASN by western blot analysis and immunofluorescence staining. Similar results as mRNA expression were obtained in protein expression (Fig. 4d; Supplementary Fig. 4a, b). Similar to the results of cell phenotype (Supplementary Fig. 3e, f), the addition of sodium acetate has no effects in both NFYA wild-type and NFYA-deficient cells (Supplementary Fig. 4c). In both NFYA wild-type and NFYA-deficient cells, Etomoxir increased CPT1A expression, and Cerulenin decreased CPT1A expression, but no difference in behavior was observed in both the presence and absence of NFYA (Supplementary Fig. 4d). These results indicate that NFYA regulates breast cancer malignant behavior by controlling de novo lipogenesis by regulating ACACA and FASN expression.

**Fig. 4.**
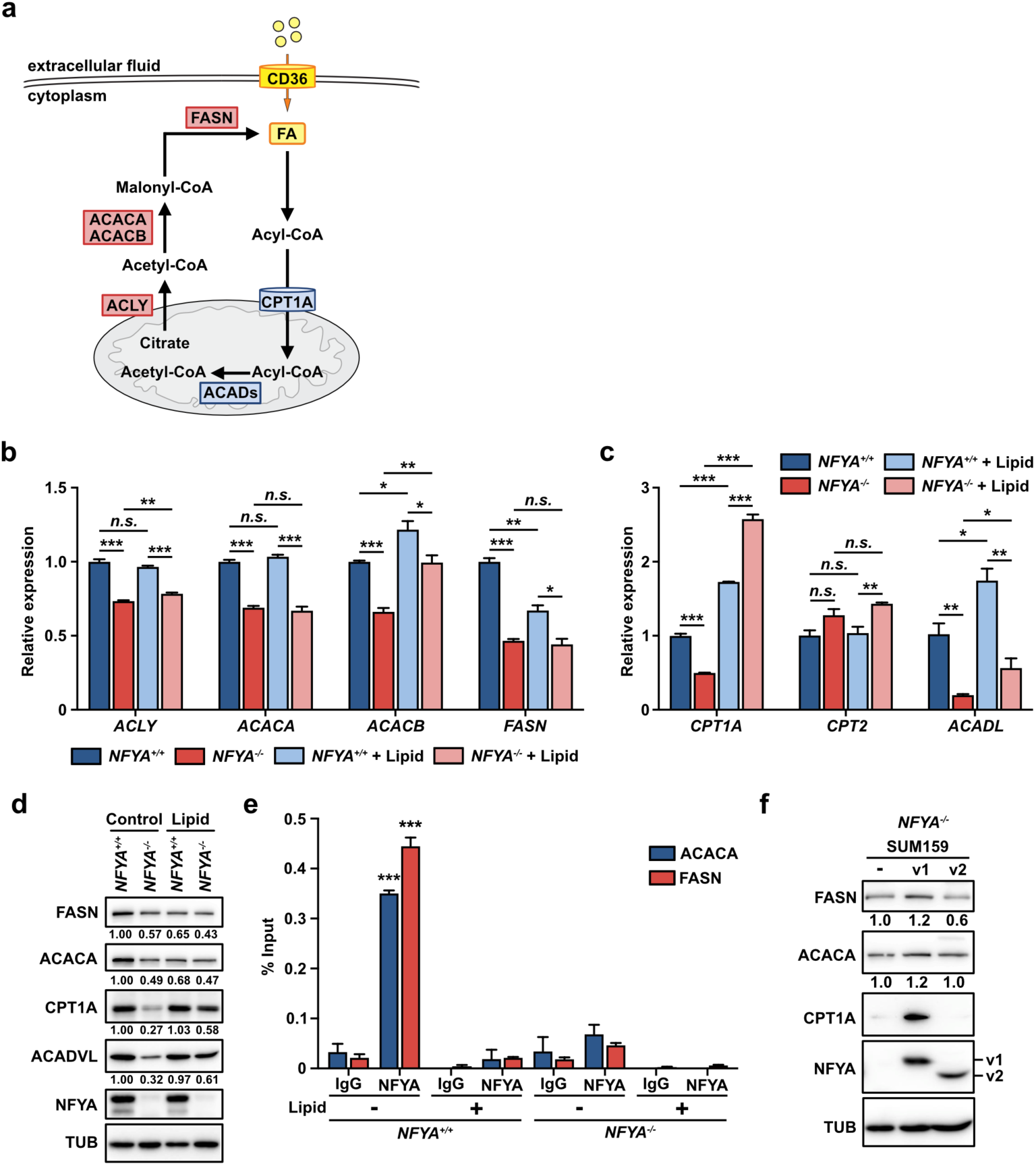
NFYA enhances lipogenesis by transcriptional activation of ACACA and FASN. **a** A diagram illustrating the reaction of lipogenesis and FAO. **b-d** qRT-PCR (**b**, **c**) and western blot analysis (**d**) of the expression levels of differentially expressed genes (**b**; Lipogenesis-related genes, **c**; FAO-related genes) in *NFYA^+/+^*and *NFYA^-/-^* SUM159 cells treated with or without lipid mixture. **e** CUT&RUN assay to evaluate the association of NFYA with the promoter of ACACA and FASN in *NFYA^+/+^* and *NFYA^-/-^* SUM159 cells treated with or without lipid mixture. Input or eluted chromatin was subjected to qRT-PCR analysis using promoter-specific primers. Data represents the % input of the immunoprecipitated chromatin for each gene. **f** Western blot analysis of the expression of FASN, ACACA, and CPT1A in NFYAv1 or NFYAv2 overexpressed NFYA-deficient SUM159 cells. All error bars represent SEM; (n.s.) not significant; (*) P<0.05; (**) P<0.01; (***) P<0.001.

We further examined whether NFYA exerts the transcriptional regulation of ACACA and FASN through direct binding to those promoter regions by CUT&RUN assay using immunoprecipitation validated antibody (Supplementary Fig. 4e). Since it is known that NFYA binds to the CCAAT box in the promoter regions of target genes^14^, we searched for the CCAAT box in the promoter regions and found ACACA at 2,825 bp upstream and FASN at 577 bp and 581 bp upstream (Supplementary Fig. 4g). Analysis of NFYA binding to the region showed that NFYA binds directly to the promoter regions of ACACA and FASN (Fig. 4e). Interestingly, adding lipid mixture to the culture medium completely abolished the binding of NFYA to the promoter regions. Furthermore, overexpression of NFYAv1, but not NFYAv2, in NFYA-deficient SUM159 cells upregulates FASN, ACACA, and CPT1A expression, supporting evidence that NFYAv1 regulates lipid metabolism via transcriptional regulation of FASN and ACACA (Fig. 4f). These results indicate that the regulation of ACACA and FASN expression by NFYA is mediated by direct binding to their promoter regions and, in addition, that in the presence of excess lipids, NFYA binding to the promoter regions is abolished, thereby regulating lipid synthesis through the downregulation of ACACA and FASN expression, as also appears in Fig. 4d. In association with these results, the fact that NFYA-deficient cells upregulated the expression of fatty acid translocase (CD36), fatty acid transporter (Supplementary Fig. 4f), suggests the presence or absence of fatty acids in the vicinity of TNBC cells alters their regulation by NFYA.

### Nfyav1 enhances tumorigenesis via the regulation of Acaca and Fasn expression in vivo

To investigate whether the regulation of lipid metabolism by NFYAv1 is involved in breast cancer tumorigenesis in vivo, we generated Nfyav1-specific knockout mice using the CRISPR/Cas9 system (Fig. 5a). We designed gRNAs on intron 2 and intron 3, introduced them into zygotes simultaneously with Cas9 protein by electroporation, and transplanted the treated embryos into pseudopregnant recipient mice to obtain founder (F0) mice. To screen Nfyav1 deficient mice, we analyzed the F0 mice by genomic DNA genotyping and sequencing analysis (Supplementary Fig. 5a, b). It has been reported that total knockout of Nfya causes embryonic lethality in early development^32^, and knockout of the lipogenic enzymes, Acly, Acaca, and Fasn, also causes embryonic lethality in early development^10–12^. Therefore, it was expected that Nfyav1 knockout mice would also show embryonic lethality, but surprisingly, Nfyav1 knockout mice were born normally (Supplementary Table 1). We crossed F0 mouse deficient in Nfyav1 with wild-type B6 mouse and used the mice obtained after the F1 generation for further analysis. The loss of Nfyav1 mRNA and protein expression in *Nfyav1^-/-^* mice was confirmed by qRT-PCR and western blot analysis in the mammary epithelial cells and MEFs (Fig. 5b, c; Supplementary Fig. 5c, d). Interestingly, *Nfyav1^-/-^* mice showed a deletion of Nfyav1 expression and a significant increase in Nfyav2 expression. Although this expressive feature is similar to that of NFYAv1-deficient cells, which suppress TNBC cell proliferation (Fig. 2g; Supplementary Fig. 2f), *Nfyav1^-/-^* mice were born at the expected Mendelian ratio, with no differences between the sexes (Fig. 5d). In addition, we explored physical characteristics in *Nfyav1^-/-^* mice and found no apparent abnormalities in mouse weight, mammary gland morphology, or survival (Fig. 5e, f; Supplementary Fig. 5e). These results suggest that NFYAv1 inhibition might be a safer therapeutic target than lipogenesis blockade by inhibiting lipogenic enzymes.

**Fig. 5.**
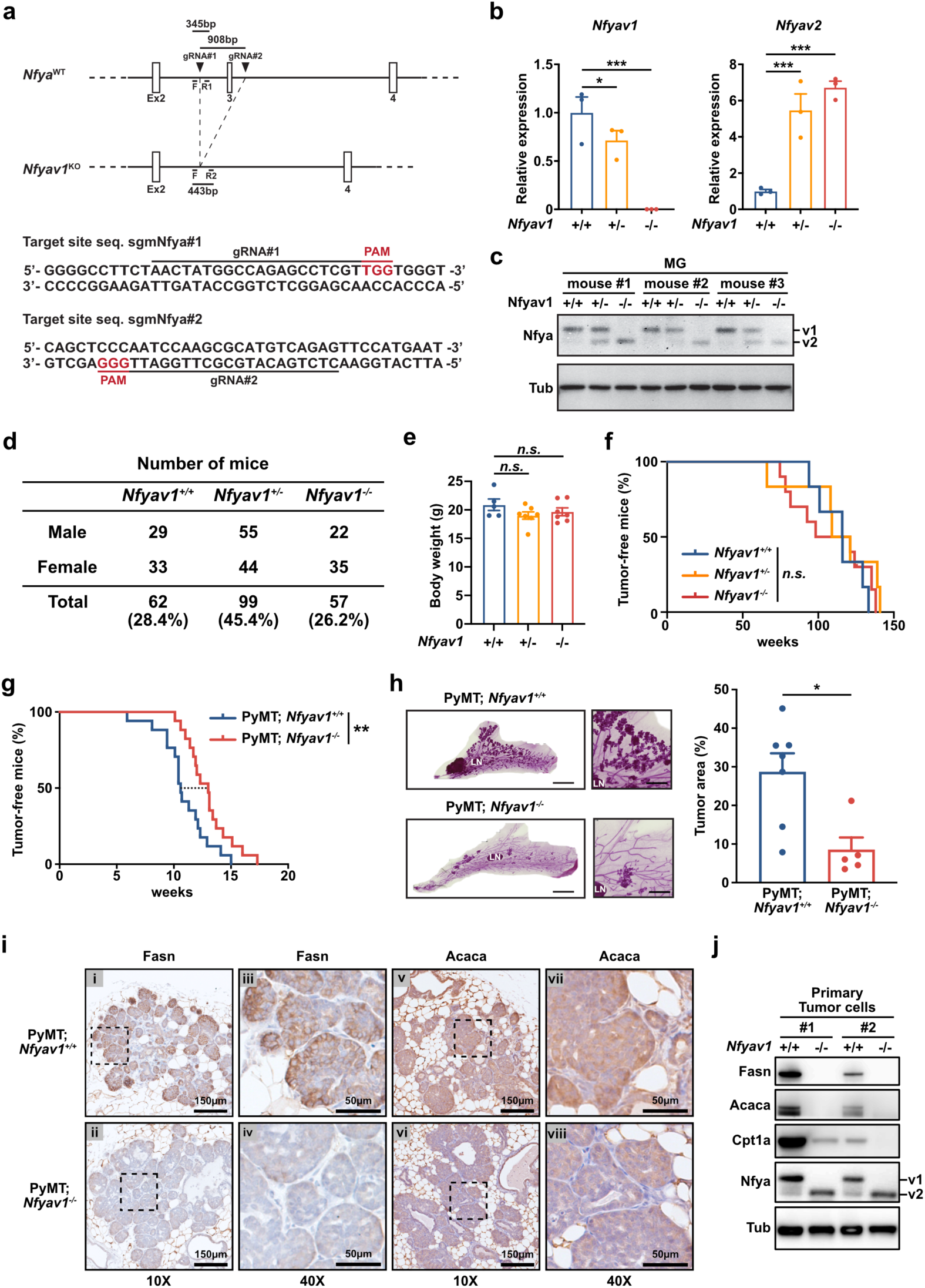
Nfyav1 enhances tumorigenesis via the regulation of Acaca and Fasn expression in vivo. **a** Diagram of endogenous *Nfya* gene structure and Nfyav1 knockout structure. Using the CRISPR/Cas9 system with two gRNAs targeted intron 2 and 3, we deleted the exon3 from genome DNA. **b, c** Confirming loss of Nfyav1 expression in mouse mammary epithelial cells. Littermate-controlled *Nfyav1^+/+^*, *Nfyav1^+/-^*, and *Nfyav1^-/-^* mice were analyzed by qRT-PCR (**b**) and western blot analysis (**c**) to qualify the expression of Nfyav1. n=3; All error bars represent SEM; (*) P<0.05; (***) P<0.001. **d** Genotypes of offspring from *Nfyav1^-/-^* mouse intercrosses. **e** Body wight of mice at 8-week-old for *Nfyav1^+/+^* (n=5), *Nfyav1^+/-^* and *Nfyav1^-/-^* (n=7). **f** Kaplan-Meier survival curve for *Nfyav1^+/+^* and *Nfyav1^+/-^* (n=6), and *Nfyav1^-/-^* (n=10). **g** Kaplan-Meier analysis of tumor-free mice for MMTV-PyMT; *Nfyav1^+/+^* (blue, n=17) and MMTV-PyMT; *Nfyav1^-/-^* (red, n=17). (**) P<0.01. **h** Representative images of whole-mount carmine alum staining of MMTV-PyMT; *Nfyav1^+/+^* or MMTV-PyMT; *Nfyav1^-/-^* mammary gland at 15-week-old. LN, lymph node. A bar graph shows the percentage of tumor area versus the total mammary fat pad in MMTV-PyMT; *Nfyav1^+/+^* (n=7) or MMTV-PyMT; *Nfyav1^-/-^* (n=5) mice. (*) P<0.05. **i** Breast cancer sections from both genotypes were stained with Fasn (panels i-iv) and Acaca (panels v-viii). **j** Protein expression analysis in PyMT induced tumors from *Nfyav1^+/+^*or *Nfyav1^-/-^* mice.

To investigate the effects of Nfyav1 on the mammary gland tumorigenesis, we generated a genetically engineered mouse model of breast cancer based on widely employed MMTV-PyMT mice^33, 34^ intercrossed with *Nfyav1^-/-^* mice. In MMTV-PyMT; *Nfyav1^+/+^* mice, mammary tumors developed as early as 41 days, and 50% of the mice developed tumors by 74 days. In contrast, in MMTV-PyMT; *Nfyav1^-/-^* mice, the first tumor was identified at 71 days, and 50% of the mice developed tumors by 91 days (Fig. 5g). Consistent with the delay of tumor occurrence, whole-mount carmine alum staining at 15 weeks showed that MMTV-PyMT; *Nfyav1^-/-^* mice possess a reduced tumor area to 29.6% of that of MMTV-PyMT; *Nfyav1^+/+^*mice (Fig. 5h; Supplementary Fig. 5f).

Immunohistochemistry with Acaca and Fasn antibodies in tumor tissues showed high expression of those genes in MMTV-PyMT; *Nfyav1^+/+^* mice. In contrast, the expression of Acaca and Fasn was strikingly decreased in the tumor tissues from MMTV-PyMT; *Nfyav1^-/-^* mice (Fig. 5i). Furthermore, we isolated breast cancer cells from these mice and analyzed the expression of those genes by western blot analysis. The results showed, similar to the immunohistochemical analysis that the expression of Acaca and Fasn was decreased in the cells isolated from *Nfyav1^-/-^* breast cancer tissues. At the same time, the expression of Cpt1a, a critical rate-limiting enzyme of FAO, was also decreased (Fig. 5j). These results indicate that loss of Nfyav1 causes a decrease in FAO by suppressing Acaca and Fasn expression in vivo, thereby suppressing breast cancer carcinogenesis, making Nfyav1 inhibition a safer and more compelling candidate for TNBC therapeutic target.

## Discussion

Here we describe the function of NFYA in the regulatory mechanism of tumor malignant behavior. Using in vivo and in vitro studies, we show that NFYAv1 strongly supports cell growth and tumorigenesis of TNBC. NFYAv1 can regulate de novo lipid synthesis and, through this regulation, can promote the malignant behavior of cancer cells. This regulation is accomplished by directly promoting ACACA and FASN transcription by NFYAv1.

Typically, the lipogenesis pathway activity is at a superficial level in normal tissues. However, high activity has been reported in TNBC to produce membrane phospholipids, signaling molecules, and energy for proliferation^5–7^ (Supplementary Fig. 3a). Therefore, the lipogenesis pathway is considered a potential therapeutic target for TNBC, for which no effective therapeutic target has been identified. In preclinical models of TNBC, inhibition of FASN alone or combination inhibition with EGFR has been reported to have a practical antitumor effect on TNBC^35–37^. However, there are still unsolved issues in the antitumor effects of lipogenesis pathway blockade.

First, FASN expression varies widely among TNBC patients, resulting in differential therapeutic effects^5, 6^. Pathological examination showed that approximately 45 % of the TNBC studied had high expression of FASN, while the remaining 55 % had low or no expression of FASN^5, 6^. It is reported that TNBC is a very heterogeneous subtype, and differences in gene expression signatures between patients are not entirely surprising^1^. Therefore, to obtain the therapeutic benefit of lipogenesis pathway blockade, it is necessary to understand lipogenesis’s regulatory mechanisms and determine which patients may benefit from this treatment. Our analysis of NFYA splicing variants’ expression in breast cancer subtypes indicates a strong correlation between the expression pattern of NFYA splicing variants and breast cancer subtypes. The Claudin^low^ breast cancer subtype, more aggressive, difficult to treat, and with poor prognosis, was characterized by a higher NFYAv1/NFYAv2 ratio.

Furthermore, we indicated that NFYAv1 promotes breast cancer malignant behavior in this subtype. It was also shown that this phenotype is due to NFYAv1 promoting transcription of lipogenic enzymes, ACACA and FASN. In addition, these phenotypes in Claudin^low^ breast cancer cells are consistent with the phenotype of delayed breast cancer development in MMTV-PyMT; *Nfyav1^-/-^* mice. In short, the NFYAv1-lipogenesis axis enhances the malignant behavior in the Claudin^low^ breast cancer subtype, and disruption of this axis inhibits its malignant behavior, indicating that the NFYA expression pattern may be helpful as a marker to determine the therapeutic efficacy of the lipogenesis pathway blockade.

The second issue is the clinical antitumor effects of lipogenesis pathway blockade. Zathy et al. showed that the effect of FASN inhibition on cell proliferation *in vitro* does not necessarily translate to the same effect on antitumor efficacy in vivo^38, 39^. In this study, we indicated that lipogenesis pathway blockade by NFYA deficiency suppresses the malignant behavior of the Claudin^low^ breast cancer cells. However, we also indicated that the addition of exogenous lipids restores the suppressed malignant behavior. Furthermore, the addition of exogenous lipids suppresses the binding of NFYA to the promoter regions of ACACA and FASN and suppresses the expression of those genes. Interestingly, NFYA deficiency increases the expression of fatty acid translocase (CD36), fatty acid transporter. These results suggest that the antitumor effect of lipogenesis pathway blockade is limited and that cancer cells overcome the antitumor effect by uptaking exogenous lipids when the lipogenesis pathway is blocked. It has been reported that cancer-associated adipocytes (CAAs), which exist in the tumor microenvironment, release free fatty acids into the tumor tissue, which are then uptaken by cancer cells to promote cancer progression ^40, 41^. In short, the reason for the limited antitumor effect of lipogenesis pathway blockade in vivo can be explained, at least in part, by the utilization of this mechanism by cancer cells. Nevertheless, several studies have confirmed the therapeutic effects of FASN inhibitor^36, 38, 42, 43^. In our in vivo mouse model, tumorigenesis is also suppressed by losing the NFYAv1- lipogenesis axis. These results suggest that not all cancer cells can overcome lipogenesis pathway blockade by uptaking exogenous lipids. Whether this difference is due to subtype, cancer progression, or the tumor microenvironment is currently unknown, and further research is needed.

Currently, therapeutic strategies that inhibit CPT1A for FAO blockade are also being tested^44^. However, FAO blockade has little effect on the production of membrane phospholipids and signaling molecules, thus limiting the antitumor effect, and lipogenesis blockade would be more advantageous. Indeed, in our study, the FASN inhibitor, Cerulenin, suppressed cellular proliferation more than the CPT1A inhibitor, ETO. It has been reported that dual-targeted therapy for prostate cancer with inhibition of de novo lipogenesis and uptaking of fatty acids potently inhibits tumor growth compared to single-targeted therapy^45^. In the future, it will be necessary to consider a combined therapeutic strategy of lipogenesis pathway, lipid uptake, and FAO blockade for curing breast cancer.

Finally, the most serious issue is safety as a therapeutic target. It has been reported that knockout mice of the lipogenic enzymes, Acly, Acaca, and Fasn, exhibit embryonic lethality in early embryogenesis^10–12^. These phenotypes raise concerns about the safety of considering the lipogenesis pathway blockade by inhibiting these genes as a therapeutic target. It has been reported that Nfya knockout mice also show embryonic lethality in early embryogenesis^32^. In this study, we generated NFYAv1-specific knockout mice born normally and did not show any apparent morphological abnormalities and early onset of diseases.

Given its safety and therapeutic efficacy, these results indicate that NFYAv1 may be a valuable and attractive therapeutic target in the treatment of TNBC. Recently, inhibitors against NF-Y have begun to be identified^46, 47^. These inhibitors bind to NF-Y and inhibit NF-Y-DNA complex formation, thereby blocking transcriptional promotion by NF-Y. However, NFYA splicing variant-specific inhibitors have not been identified because the mechanism that induces transcription of different target genes among NFYA splicing variants is still unclear. Further studies would be necessary to elucidate the details of the mechanism of transcriptional regulation by NFYA splicing variants.

This study reveals an essential role for NFYA in regulating lipid metabolism and the regulatory mechanisms of tumor malignant behavior by the NFYA-lipogenesis axis, demonstrating the importance of the lipid metabolic pathway as a therapeutic target for TNBC.

## Methods

### Generation of Nfyav1 knockout mice

Electroporation introduced two single guide RNAs (sgRNA) targeting the intron 2 and 3 of Nfya (sgmNfya#1: 5’-AACTATGGCCAGAGCCTCGT-3’ and sgmNfya#2: 5’-CTCTGACATGCGCTTGGATT-3’) and human codon-optimized Cas9 into zygotes obtained from C57BL/6J to remove exon 3 of Nfya (Fig. 5a). Treated zygotes were cultured overnight to the 2-cell stage and then transferred to pseudopregnant recipient mice. Founder mice were genotyped using primers (primer F: 5’-GAGTCCCAAGCCACTGATGA-3’, primer R1: 5’-ACCATGGATGAAGGAACTAGCC-3’, and primer R2: 5’-TCCTGCCTCCATATCCCAAC-3’) (Fig. 5a). To generate a PyMT-induced mouse breast cancer model, we obtained FVB/N-Tg(MMTV-PyVT)634Mul/J from The Jackson Laboratory (no. 002374) and backcrossed with C57BL/6N females. After six generations, MMTV-PyMT mice were crossed with *Nfyav1^-/-^* mice to generate MMTV-PyMT; *Nfyav1^-/-^* mice.

### Animal experiments

All mouse experiments were performed in accordance with Kyoto University, Kanazawa University, and Okayama University Institutional Animal Care and Use Committee approved protocol (Med Kyo13605, AP-153426, OKU-2018654, OKU-2018906). For orthotopic tumor growth assay, 1 × 10^6^ cells were resuspended in 30 µl culture media containing 50 % Matrigel (Corning, no. 356230) and injected into the mammary fat pad of 6-week-old NOD-SCID female mice. Mice were sacrificed 50 days later to determine the tumor weight.

### Isolation of mouse tumor cells

Primary mouse breast tumor cells were isolated from individual breast tumors. Following harvest, tumors were cut into small pieces and digested for 1.5 hours at 37 °C in 5 ml of DMEM/F12 containing 2 mg/ml of collagenase type I (WAKO, no. 037-17603) and 100 U/ml of hyaluronidase (Sigma, no. H3506). Following digestion, trypsin/EDTA was added, and the mixture was incubated for 5 min at 37 °C. Digested tumor samples were then pressed through 40-µm cell strainers. Finally, samples were centrifuged at 1,000 rpm for 5 min at room temperature, resuspended in DMEM supplemented with 10 % FBS and 1 % penicillin and streptomycin, and plated on collagen type I coated 100 mm dishes (IWAKI, no. 4020-010-MYP).

### qRT-PCR analysis

According to the manufacturer’s instruction, total RNA was extracted from cells and mammary glands using TRIzol reagent (Thermo Fisher Scientific, no. 15596018). For qRT-PCR analysis, cDNA was synthesized from total RNA using a High-Capacity RNA-to-cDNA Kit (Applied Biosystems, no. 4387406). mRNA expression levels were determined by qRT-PCR with KAPA SYBR FAST qPCR Master Mix Kit (Kapa Biosystems, no. KK4610). Relative expression levels were normalized to human *ACTB* or mouse *Actb*. The primers used here are shown in Supplementary Table 2.

### Western blot analysis

Total cellular extracts resolved by SDS-PAGE were transferred to PVDF membranes. Western blot was performed in TBST (100 mM Tris-HCl at pH7.5, 150 mM NaCl, 0.05 % Tween-20) containing 5 % Blocking One (nacalai tesque, no. 03953-95). Immunoreactive protein bands were visualized using SuperSignal West Pico PLUS Chemiluminescent Substrate (Thermo Scientific, no. 34577).

### Cell culture

All human breast cancer cells, NMuMG cells, and 293T cells were obtained from American Type Culture Collection. HMLE cells were obtained from R. Weinberg (MIT, USA). MDAMB231, NMuMG, and 293T cells were maintained in DMEM supplemented with 10 % FBS. MCF7 cells were maintained in DMEM supplemented with 10 % FBS and 10 μg/ml insulin.

BT474, HCC1143, BT549, and MDAMB468 cells were maintained in RPMI1640 supplemented with 10 % FBS. SUM149 and SUM159 cells were maintained in Ham’s F12 supplemented with 5 % FBS, 10 mM HEPES, 1 µg/ml Hydrocortisone, and 5 μg/ml Insulin. MCF10A and MCF12A cells were maintained in MEGM supplemented 100 ng/ml cholera toxin. HMLE cells were maintained in MEGM bullet kit (Lonza, no. CC-3150). All media were supplemented with 1 % penicillin and streptomycin.

### CRISPR/Cas9 system

CRISPR/Cas9 system was used to knockout the NFYA gene in SUM159, MCF7, and HMLE cells. NFYA and NFYAv1-specific gRNAs were determined using the candidates provided by GPP sgRNA Designer (https://portals.broadinstitute.org/gpp/public/analysis-tools/sgrna-design) and cloned into px459 vector (Addgene, no. 48139) for NFYA knockout and lentiCRISPRv2 vector (Addgene, no. 52961) for NFYAv1-specific knockout. Sequence for human NFYA sgRNA #1: GCCTTACCAGACAATTAACC, human NFYA sgRNA#2: GAGCAGATTGTTGTCCAGGC, human NFYAv1 sgRNA#1: GCCCAGGTGGCATCCGCCTC, human NFYAv1 sgRNA#2: GGCCTGAGGCGGATGCCACC.

### Generation of retrovirus and lentivirus

PCR products were then cloned into pMSCV-puro, pTRE3G-blast, or lentiCRISPRv2. For retrovirus preparation, 293T cells were transfected with 300 ng of pMSCV-puro-NFYAv1, -NFYAv2, - NFYAv1YA29mt, or -NFYAv2YA29mt together with 300 ng of packaging plasmid, pCL-Ampho. For lentivirus preparation, 293T cells were transfected with 300 ng of pTRE3G-blast-SNAIL or lentiCRISPRv2-gRNA together with packaging plasmids (300 ng of pMDLg/pRRE, 150 ng of pMD2g, and 150 ng of pRSVRev). Cells were infected using the filtrated culture supernatant from 293T cells in the presence of 8 µg/ml polybrene.

### Cell proliferation

For direct cell counting experiments, tumor cells were plated in triplicate at 0.2 × 10^4^ cells per well of a 24-well plate. At indicated days, cells were trypsinized and counted.

For crystal violet staining experiments, tumor cells were plated 0.5 × 10^4^ cells per well of a 24-well plate. At indicated days, cells were washed with PBS (-) and stained with 0.5 % crystal violet solution for 20 min. The wells were washed with water, dried, and incubated with methanol for 20 min. The amount of dye reflecting the cell number was measured at 590 nm.

### Sphere formation assay

Single-cell suspensions of cell lines were suspended at a density of 2,000 cells/ml in DMEM/Ham’s F-12 (nacalai tesque, no. 11581-15) containing 20 ng/ml bFGF (Peprotech, no. 100-18B), 20 ng/ml EGF (Wako, no. 059-07873), 1 × B27 (Thermo Scientific, no. 17504044), and 1 % methylcellulose 400 (Wako, no. 132-05055) into 24-well ultra-low attachment plates (Corning, no. 3473). Spheres were counted after 7 days.

### Bodipy staining

Cells were cultured on 8 wells chamber slides with or without 2 % lipid mixture for 3 days and fixed with 4 % paraformaldehyde for 20 min at room temperature. After washing, cells were incubated with 3.8 µM of BODIPY 493/503 (Thermo Fisher Scientific, no. D3922) for 30 min at room temperature, then mounted with Vectashield hard-set mounting medium with DAPI (Vector laboratories, no. H-1500).

### Oxygen Consumption Rate (OCR) measurement

5 × 10^4^ cells were plated on XF24 cell culture plate (Agilent, no. 100777-004) with DMEM supplemented with 1 % FBS, 0.5 mM glucose, 1 mM Glutamax, 0.5 mM Carnitine, 10 mM HEPES, 1 µg/ml Hydrocortisone, and 5 µg/ml Insulin. For measuring OCR in response to lipid stimulation, cells were cultured with 33 µM Palmitate-BSA or free fatty acid reduced-BSA in Krebs-Henseleit buffer supplemented with 2.5 mM Glucose, 0.5 mM Carnitine, 5 mM HEPES, 1 µg/ml Hydrocortisone, and 5 µg/ml Insulin at 37 °C for 1 hour without CO_2_ control. 1 µM Oligomycin A (Cell Signaling Technology, no. 9996) was injected to inhibit ATPase V. Maximal OCR was induced by exposing cells to mitochondrial uncoupler, 2 µM FCCP (Sigma-Aldrich, no. C2920). 1 µM Antimycin A (Sigma-Aldrich, no. A8674) and 1 µM rotenone (Sigma-Aldrich, no. R8875) were added to disrupt all mitochondria-dependent respiration. XF24 was used to measure OCR under a 3 min period protocol, followed by 2 min mixing and 3 min incubation.

### CUT & RUN analysis

CUT & RUN experiments were performed using 3 × 10^5^ cells with the CUT&RUN assay kit (CST, no. 86652). In brief, cells were washed, bound to activated Concanavalin A magnetic beads, and permeabilized with antibody binding buffer containing Digitonin. The bead-cell complex was incubated overnight with 0.7 µg of NFYA monoclonal antibody at 4 °C. The bead-cell complex was washed with Digitonin buffer and incubated pAG-MNase solution for 1 hr at 4°C. After washing with Digitonin buffer, 2 mM calcium chloride was added to activate pAG-MNase and incubated for 30 min at 4 °C. After incubation, the reaction was stopped with stop buffer. DNA fragments were released by incubation for 10 min at 37 °C and purified with Fast Gene Gel/PCR extraction kit (Nippon genetics, no. FG-91202). The DNA fragments were quantified by qRT-PCR analysis. The primers used here are shown in Supplementary Table S2.

### Histology and immunohistochemistry (IHC)

For IHC, paraffin sections were deparaffinized, dehydrated, and subjected to heat-induced antigen retrieval in a pressure cooker using 10 × G-Active pH6 (Geno Staff, no. ARSC6-01). Slides were incubated for 10 min with 3 % H_2_O_2_, blocked for 1 hour with BlockingOne Histo (nacalai tesque, no. 06349), and incubated with primary antibody overnight at 4 °C in Can Get Signal immunostain solution A (TOYOBO, no. NKB-501). Signals were enhanced using Vectastain ABC Elite kit (Vector Laboratories, no. PK-6101) and visualized by ImmPACT DAB (Vector Laboratories, no. SK-4105) and counterstaining with Mayer’s Hematoxylin (Wako, no. 131-09665).

For immunofluorescence staining, paraffin sections were deparaffinized, dehydrated, and subjected to heat-induced antigen retrieval in a pressure cooker using 10 × G-Active pH6. Slides were blocked for 1 hour with BlockingOne Histo and incubated with primary antibody overnight at 4 °C in 5 % BlockingOne Histo in PBS (-). The next day, the slides were labeled with Alexa Fluor 568 (Thermo Fisher Scientific, no. A-11036) or Alexa Fluor 488 (Thermo Fisher Scientific, no. A-11011) and mounted the slides with Vectashield hard-set mounting medium with DAPI (Vector laboratories, no. H-1500).

For whole-mount carmine-alum staining, mammary fat pads, including mammary gland, were fixed on glass slides with fixation buffer (25 % acetic acid, 75 % ethanol) overnight. Tissues were hydrated and stained with carmine alum solution (Stemcell technologies, no. 07070). Tissues were dehydrated, cleared in Histoclear (National Diagnostics, no. HS-200), and mounted with Permount (Falma, no. SP15-100-1).

### Antibodies

The following monoclonal (mAb) and polyclonal (pAb) primary antibodies were used for western blot and IHC: NFYA pAb (1:500; Santa Cruz, no. sc-10779), NFYA mAb (1:500; Santa Cruz, no. sc-17753), E-Cadherin mAb (1:1000; BD, no. 610181), Vimentin mAb (1:1000; BD, no. 550513), Flag mAb (1:1000; Sigma, no. F3165), FASN Rabbit mAb (1:1000 for western blot, 1:100 for IHC; CST, no. 3180), ACACA Rabbit mAb (1:1000 for western blot, 1:100 for IHC; CST, no. 3676), CPT1A Rabbit mAb (1:1000; abcam, no. Ab234111), ACADVL mAb (1:1000; Santa Cruz, no. sc-376239), CD36 pAb (1:500; Santa Cruz, no. sc-9154), Keratin 14 pAb (1:500; BioLegend, no. 905304), Keratin 8 mAb (1:500; BioLegend, no. 904804), and α-Tubulin mAb (1:2000; Sigma, no T5168).

### Reagents

The following reagents were used: Forskolin (10 µM; Wako, no. 067-02191), Lipid mixture 1 (2 %; Sigma, no. L0288), Etomoxir (3 µM; abcam, no. Ab254445), and Cerulenin (20 µM; Biolinks, no. BLK-0380).

### Statistical analysis

Data are presented as the mean ± SEM. Statistical significance of the difference between experimental groups was assessed using unpaired two-tailed Student’s t-test using GraphPad Prism 9 software. P values of < 0.05 were considered significant.

## Acknowledgments

We thank members of Medical Protein Engineering Laboratory, Okayama University, Division of Oncology and Molecular Biology, Kanazawa University, and DSK project, Kyoto University for their help and input. Particularly, we thank Mr. R. Ito, Mr. H. Sakai, and Dr. S. Ikeda for technical assistance and stimulating discussion. This work was supported by Grant-in-Aid for Scientific Research 15K18407 (to N.O.), Grant-in-Aid for Scientific Research 19K07640 (to N.O.), JSPS KAKENHI Grant JP16H06276 (AdAMS) (to N.O.), Cancer Research Institute of Kanazawa University (to N.O.), Takeda Science Foundation (to N.O.), Wesco Scientific Promotion Foundation (to N.O.), and The Okayama Foundation for Science and Technology (to N.O.).

## Author contributions

N.O. conceived the project and designed experiments. N.O., C.U., M.S., and G.T. performed experiments and analyzed data. S.K. and C.T. performed metabolic analysis and provided intellectual input on metabolic mechanisms. H.M. provided technical help to generate mice. K.Y. performed experiments and provided intellectual input on breast cancer. N.O. and C.T. wrote and edited the manuscript. All authors provided comments on the manuscript and gave final approval.

## Additional information

Supplementary information can be found in a separate file.

## Ethics declarations

Competing of interests

The authors have declared no competing interests.

