## Supplementary Figures for "NFYA promotes the malignant behavior of triple-negative breast cancer through the regulation of lipid metabolism"

**Okada *et al.* Supplementary Table 1**

**a**

| Sperm | Egg | No. electropolated | No. survived | No. 2-cell | 2-cell % | No. transferred (female mice) | No. birth | Birth % |
|---|---|---|---|---|---|---|---|---|
| C57BL/6J | C57BL/6J | 50 | 50 | 50 | 100 % | 50 (3) | 27 | 54.0 % |

**b**

Number of founder (F0) mice

| | *Nfyav1*$^{+/+}$ | *Nfyav1*$^{+/-}$ | *Nfyav1*$^{-/-}$ | N.D. |
|---|---|---|---|---|
| Male | 6 | 2 | 3 | 2 |
| Female | 4 | 0 | 4 | 2 |
| Total | 10 | 2 | 7 | 4 |

N.D.; not determined

**Supplementary Table 1** Generation of Nfyav1 knockout mice. **a** Table shows the information on mutation rates and production statistics following the introduction of sgRNAs and Cas9 into zygotes. **b** Genotypes of F0 mice.

**Okada *et al.* Supplementary Table 2**

| **qRT-PCR primer sequences** | |
|---|---|
| mouse Actb forward primer | 5'- GATCTGGCACCACACCTTCT -3' |
| mouse Actb reverse primer | 5'- GGGGTGTTGAAGGTCTCAAA -3' |
| mouse Nfyav1 forward primer | 5'- AAGTCCAGACCCTCCAGGTAGT -3' |
| mouse Nfyav1 reverse primer | 5'- GATGGGTTGGCCTGTTGAT -3' |
| mouse Nfyav2 forward primer | 5'- GCCATGGAGCAGTATACGACA -3' |
| mouse Nfyav2 reverse primer | 5'- CCTGGACCTGCTGCTGAA -3' |
| mouse Acly forward primer | 5'- GAAGGGAGTGACCATCATTGG -3' |
| mouse Acly reverse primer | 5'- GTTGTCCAGCATTCCACCAG -3' |
| mouse Acaca forward primer | 5'- ACCGCCAGCTTAAGGACAAC -3' |
| mouse Acaca reverse primer | 5'- GTTGAGGTTGGAGGCAAAGG -3' |
| mouse Fasn forward primer | 5'- ATAAGCCCAAGGCCAAGTACC -3' |
| mouse Fasn reverse primer | 5'- AGACACCTTCCCGTCACACA -3' |
| human ACTB forward primer | 5'- ACCAACTGGGACGACATGGAGAAA -3' |
| human ACTB reverse primer | 5'- TAGCACAGCCTGGATAGCAACGTA -3' |
| human NFYAv1 forward primer | 5'- CCAGACCCTCCAGGTAGT -3' |
| human NFYAv1 reverse primer | 5'- GCAAACCCTGTGTTCCAGA -3' |
| human NFYAv2 forward primer | 5'- ATTCAGCAGCAGGTCCAAGG -3' |
| human NFYAv2 reverse primer | 5'- GTTGGCCAGTTGATGTGATTAG -3' |
| human CDH1 forward primer | 5'- TGAAGGTGACAGAGCCTCTGGAT -3' |
| human CDH1 reverse primer | 5'- TGGGTGAATTCGGGCTTGTT -3' |
| human VIM forward primer | 5'- CTGCCAACCGGAACAATGA -3' |
| human VIM reverse primer | 5'- GGTACTCAGTGGACTCCTGCTTT -3' |
| human ACLY forward primer | 5'- TGTTGGCGTGGATGAGAAAC -3' |
| human ACLY reverse primer | 5'- AGAGGCCGGAGATAAAACTGG -3' |
| human ACACA forward primer | 5'- ACCGCCAGCTTAAGGACAAC -3' |
| human ACACA reverse primer | 5'- AGTGGTTGAGGTTGGAGGAGA -3' |
| human ACACB forward primer | 5'- GGACTTCCCGATCCTTTTCAG -3' |
| human ACACB reverse primer | 5'- CGACCAAACAGAGACACAGCA -3' |
| human FASN forward primer | 5'- TCCACCAGCAACATCAGCTC -3' |
| human FASN reverse primer | 5'- TTCTCCAGCAAGCCATCTCTC -3' |
| human CPT1A forward primer | 5'- TGATGACGGCTATGGTGTGTC -3' |
| human CPT1A reverse primer | 5'- TTGCTTCTTTCAGGTGCCTTC -3' |
| human CPT2 forward primer | 5'- CAGCAGATGATGGTTGAGTGC -3' |
| human CPT2 reverse primer | 5'- CCGCAGAGCAAACAAGTGTC -3' |
| human ACADL forward primer | 5'- TGGCAAAACAGTTGCTCACC -3' |
| human ACADL reverse primer | 5'- CACAAATGCTCGGGTTACACA -3' |

| **Primer sequences for ChIP-qRT-PCR** | |
|---|---|
| human ACACA forward primer | 5'- CTCCCGGGTTCAAGCGAGTC -3' |
| human ACACA reverse primer | 5'- CGCGCCTGTAATCCCAGCTAA -3' |
| human FASN forward primer | 5'- TCACCCTATCGCCTAGCAAC -3' |
| human FASN reverse primer | 5'- GCAGCAGCAACCAATCG -3' |

**Supplementary Table 2** Primer sequences for qRT-PCR analysis.

**Okada *et al.* Supplementary Fig. 1**

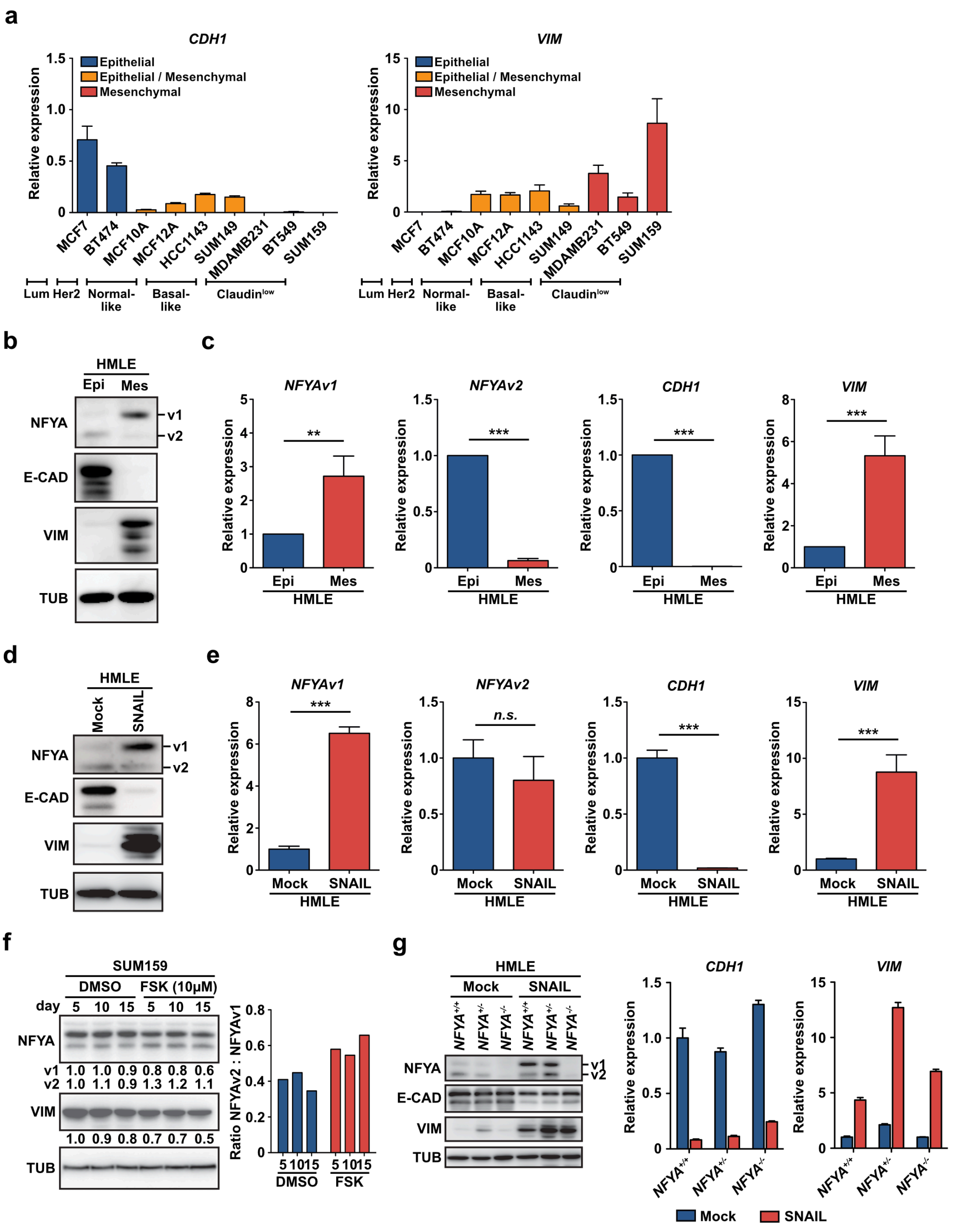

**Supplementary Fig. 1** NFYA switches the expression of alternative splicing variants during EMT progression. **a** qRT-PCR analysis of epithelial marker gene (*CDH1,* left panel) and mesenchymal marker gene (*VIM,* right panel) mRNA levels in various breast cancer cell lines. **b, c** Western blot (**b**) and qRT-PCR analysis (**c**) of NFYAv1 and NFYAv2 in HMLE-Epi and HMLE-Mes cells. E-CAD/CDH1 and VIM are markers for epithelial and mesenchymal cells, respectively. **d, e** Western blot (**d**) and qRT-PCR analysis (**e**) of NFYAv1 and NFYAv2 in HMLE cells overexpressed SNAIL to induce EMT. E-CAD/CDH1 and VIM are markers for epithelial and mesenchymal cells, respectively. **f** MET induced by 10 μM of Forskolin (FSK) treatment in SUM159 cells induced reverse switching from NFYAv1 to NFYAv2 expression. The ratio of each band is relative to the amount of each protein in DMSO treatment for 5 days. The graph shows the ratio of NFYAv2/NFYAv1. **g** Western blot and qRT-PCR analysis in $NFYA^{+/+}$, $NFYA^{+/-}$, and $NFYA^{-/-}$ HMLE cells with or without SNAIL overexpression. All error bars represent SEM; (*n.s.*) not significant; (**) P<0.01; (***) P<0.001.

**Okada *et al.* Supplementary Fig. 2**

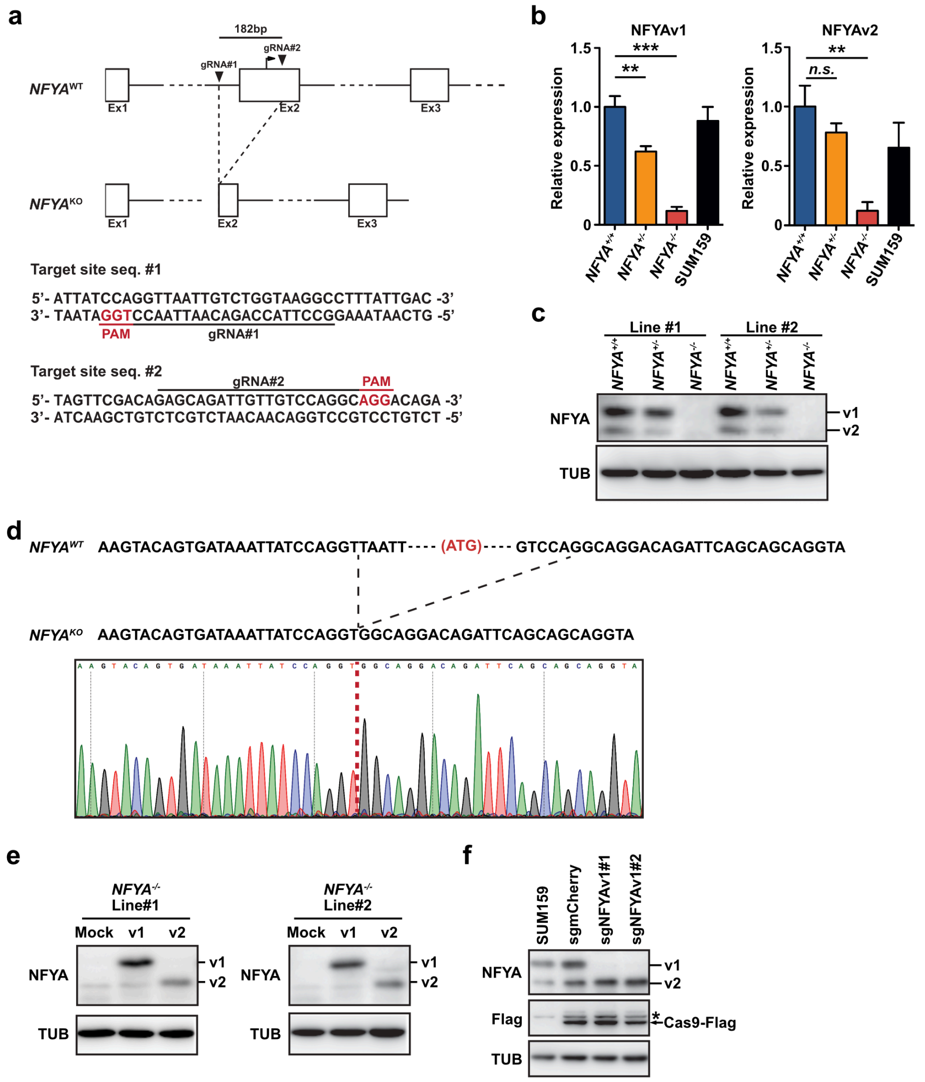

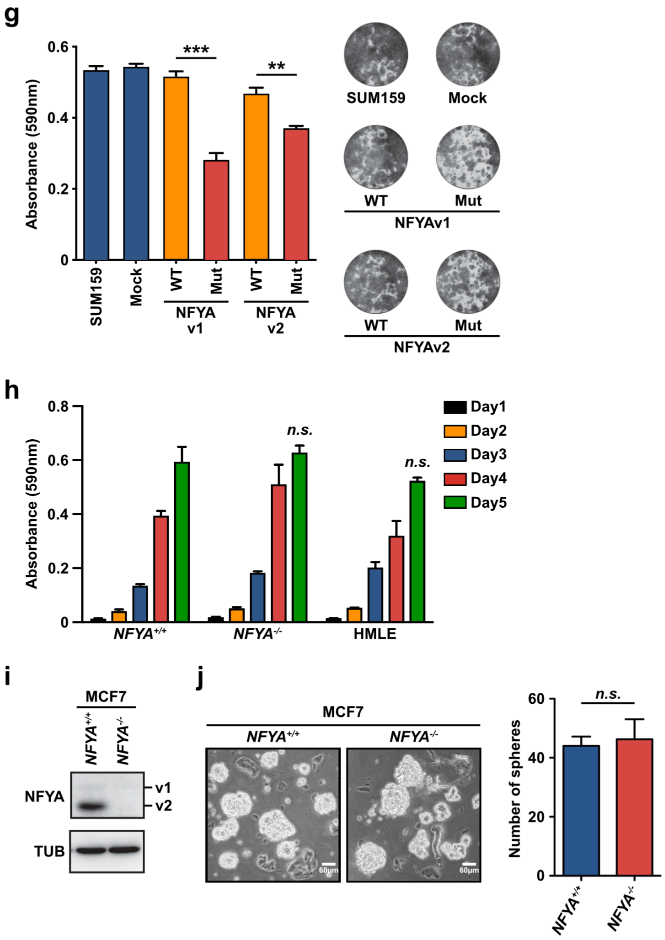

**Supplementary Fig. 2** NFYAv1 deficiency significantly inhibits tumor cell growth and tumorigenesis in TNBCs. **a** Schematic diagram of two sgRNA target sites located on both sides of the start codon and sequences of the selected sites for editing *NFYA*. Protospacer-adjacent motif (PAM) sequences are highlighted in red. **b, c** qRT-PCR (**b**) and western blot analysis (**c**) to validate the deletion of *NFYA* gene expression generated by CRISPR/Cas9 in SUM159 cells. **d** The result of sequence analysis of *NFYA*$^{-/-}$ SUM159 cells manipulated by CRISPR/Cas9. *NFYA*$^{-/-}$ SUM159 cells deleted the start codon, ATG. **e** The validation of overexpression of NFYA variants in *NFYA*$^{-/-}$ SUM159 cells. **f** *NFYAv1* specific knockout was validated by western blot analysis. Asterisk indicates nonspecific bands. sgNFYAv1#1 and sgNFYAv1#2 target different sequences. sgmCherry used as a control targeted the sequence of mCherry. **g** Quantification and representative pictures of 0.5 % crystal violet staining of SUM159 cells overexpressed WT and dominant-negative mutants of both variants of NFYA. **h** A bar graph shows the quantification of 0.5 % crystal violet staining of *NFYA*$^{+/+}$ and *NFYA*$^{-/-}$ HMLE cells. **i** Western blot analysis to validate the deletion of *NFYA* gene expression generated by CRISPR/Cas9 in MCF7 cells. **j** Representative images of sphere formation by *NFYA*$^{+/+}$ and *NFYA*$^{-/-}$ MCF7 cells. A bar graph shows the number of spheres larger than 60 μm in each group. All error bars represent SEM; (*n.s.*) not significant; (**) P<0.01; (***) P<0.001.

Okada *et al.* Supplementary Fig. 3

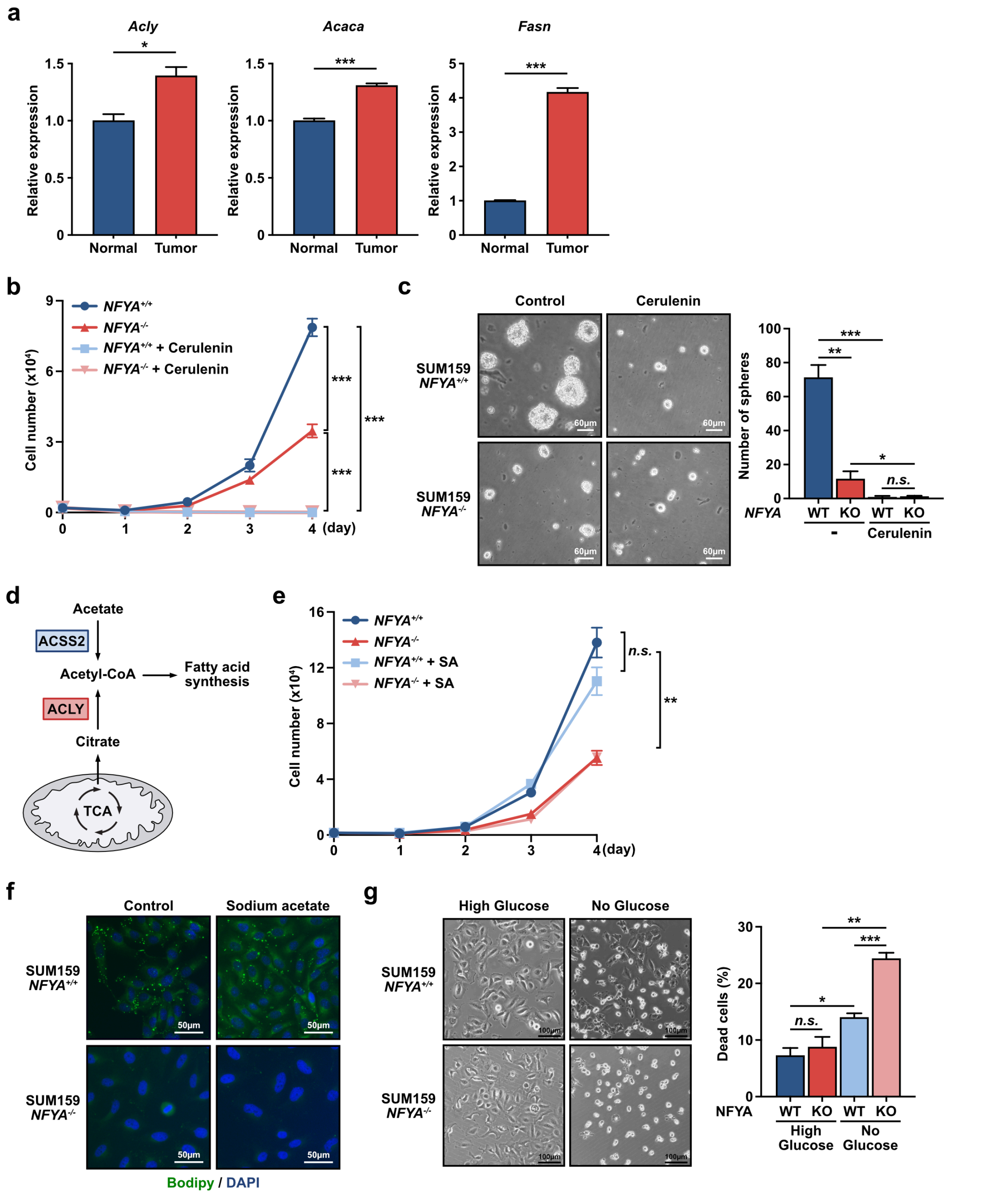

**Supplementary Fig. 3** NFYA regulates lipid metabolism for malignant behavior of TNBCs. **a** qRT-PCR analysis of *Acly*, *Acaca*, and *Fasn* in breast cancer cells (primary cells isolated from mouse breast cancer tissue) compared with non-transformed mouse mammary epithelial cells (NMuMG cells). **b** Cumulative population of cells was measured for 4 consecutive days in *NFYA*$^{+/+}$ and *NFYA*$^{-/-}$ SUM159 cells with or without the addition of Cerelenin. **c** Representative images of sphere formation by *NFYA*$^{+/+}$ and *NFYA*$^{-/-}$ SUM159 cells with or without the addition of Cerulenin. A bar graph shows the number of spheres larger than 60 µm in each group. **d** A schematic diagram depicting two different pathways for acetyl-CoA synthesis. **e** Cumulative population of cells was measured for 4 consecutive days in *NFYA*$^{+/+}$ and *NFYA*$^{-/-}$ SUM159 cells treated with or without sodium acetate. **f** Representative fluorescence images of lipid droplet (green) detected with Bodipy 493/503 and nucleus (blue) detected with DAPI in *NFYA*$^{+/+}$ and *NFYA*$^{-/-}$ SUM159 cells treated with or without sodium acetate. **g** Representative images of *NFYA*$^{+/+}$ and *NFYA*$^{-/-}$ SUM159 cells with or without glucose deprivation for 3 hours. A bar graph shows the percentage of dead cells in each group. All error bars represent SEM; (*n.s.*) not significant; (*) P<0.05; (**) P<0.01; (***) P<0.001.

Okada *et al.* Supplementary Fig. 4

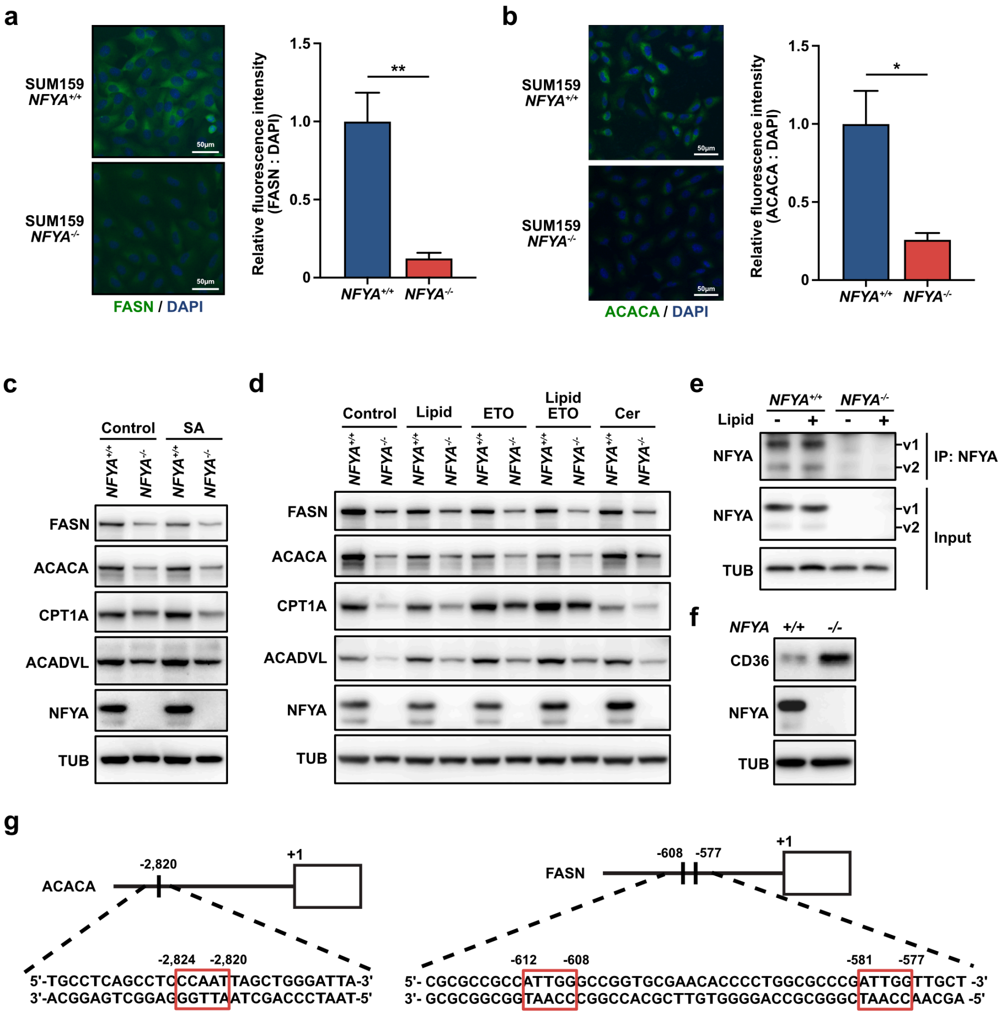

**Supplementary Fig. 4** NFYA enhances lipogenesis by transcriptional activation of ACACA and FASN. **a, b** *NFYA*$^{+/+}$ and *NFYA*$^{-/-}$ SUM159 cells were immunostained with anti-FASN (**a**, green) or anti-ACACA (**b**, green) antibodies and counterstained with DAPI for DNA (blue). A bar graph shows the relative fluorescence intensity of FASN or ACACA staining to DAPI. **c** The western blot analysis of lipogenesis and FAO related genes in *NFYA*$^{+/+}$ and *NFYA*$^{-/-}$ SUM159 cells treated with or without sodium acetate (1 mM for 3 days). **d** The western blot analysis of lipogenesis and FAO related genes in *NFYA*$^{+/+}$ and *NFYA*$^{-/-}$ SUM159 cells treated with or without lipid mixture, Etomoxir (ETO), both, and Cerulenin. **e** The western blot analysis validated immunoprecipitation of NFYA by using the anti-NFYA antibody in *NFYA*$^{+/+}$ and *NFYA*$^{-/-}$ SUM159 cells. **f** The western blot analysis of fatty acid translocase (CD36). **g** The location of NFYA binding sites on ACACA and FASN promoter.

**Okada *et al.* Supplementary Fig. 5**

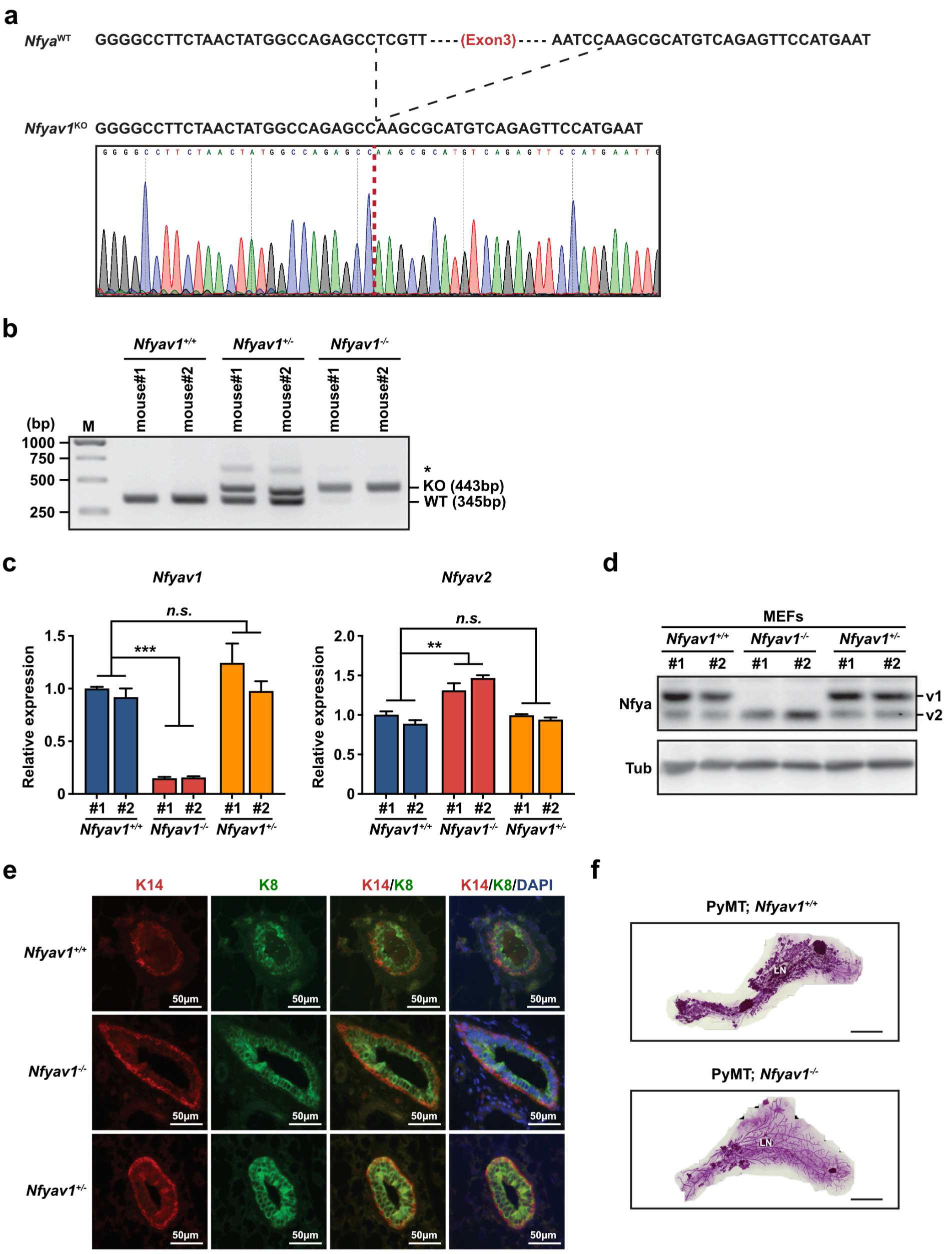

**Supplementary Fig. 5** Nfyav1 enhances tumorigenesis via the regulation of Acaca and Fasn expression *in vivo*. **a** The sequence analysis of *Nfyav1*$^{-/-}$ mouse manipulated by CRISPR/Cas9. **b** PCR analysis of genomic DNA from tails of *Nfyav1*$^{+/+}$, *Nfyav1*$^{+/-}$, and *Nfyav1*$^{-/-}$ mice using primers shown in Fig. 5A.

Amplification products correspond to WT and KO alleles (345bp and 443bp, respectively). Asterisk indicates nonspecific bands. **c, d** Confirming loss of Nfyav1 expression in *Nfyav1*$^{-/-}$ MEFs. Littermate-controlled *Nfyav1*$^{+/+}$, *Nfyav1*$^{+/-}$, and *Nfyav1*$^{-/-}$ MEFs were analyzed by qRT-PCR (**c**) and western blot analysis (**d**). Error bars represent SEM; (*n.s.*) not significant; (**) $P<0.01$; (***) $P<0.001$. **e** Immunofluorescence analysis of the K14 (red), K8 (green), and DAPI (blue) in *Nfyav1*$^{+/+}$, *Nfyav1*$^{+/-}$, and *Nfyav1*$^{-/-}$ mammary glands. **f** Representative images of whole-mount carmine alum staining of MMTV-PyMT; *Nfyav1*$^{+/+}$ or MMTV-PyMT; *Nfyav1*$^{-/-}$ mammary gland at 15-week-old. LN, lymph node.

Okada *et al.* Supplementary Fig. 6

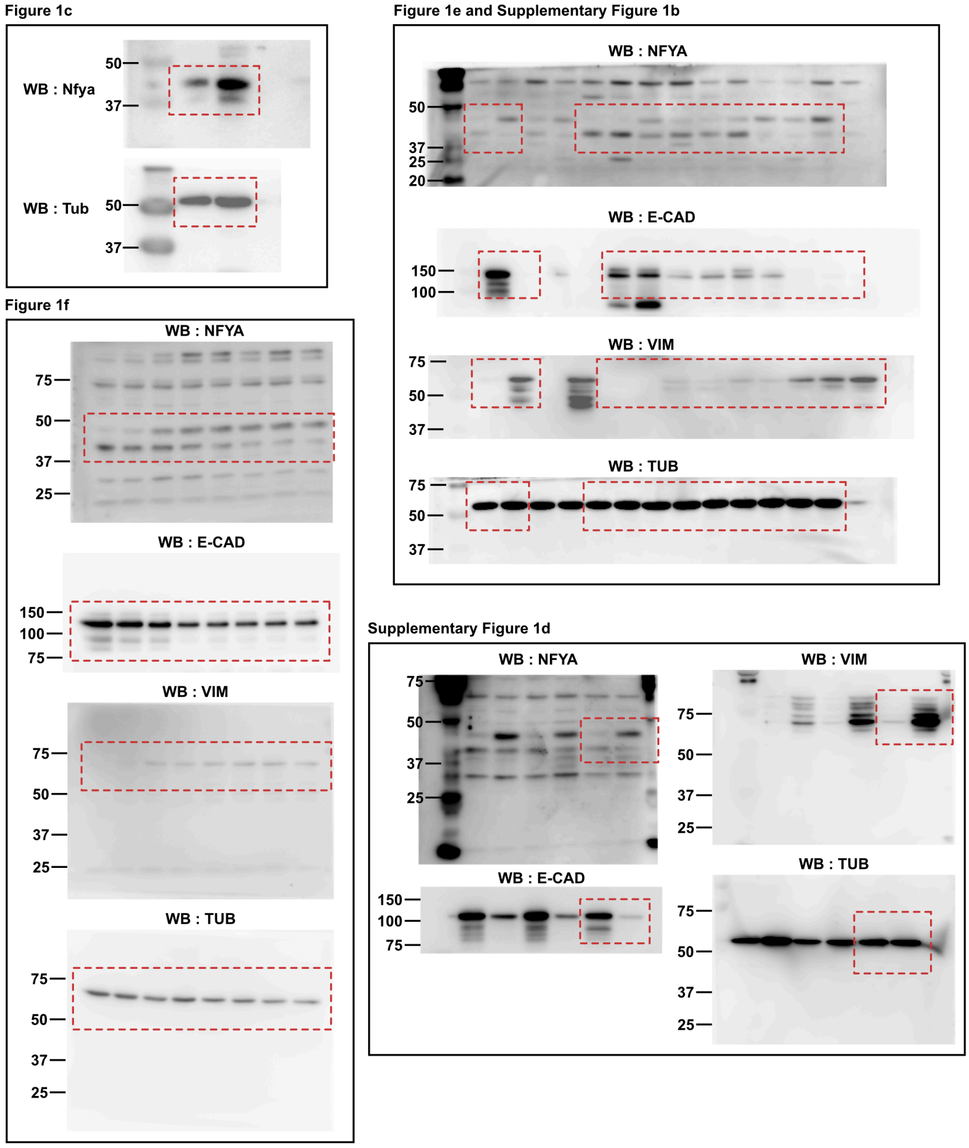

Supplementary Figure 1f

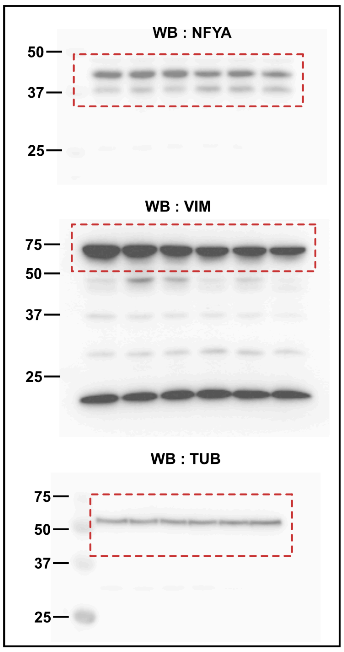

Supplementary Figure 1g

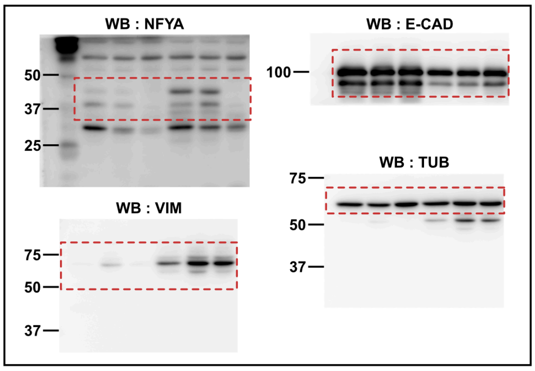

Supplementary Figure 2c

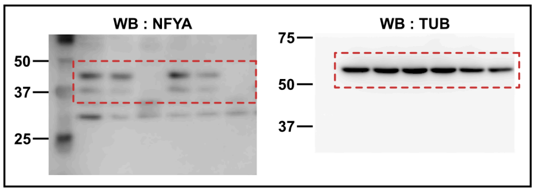

Supplementary Figure 2e

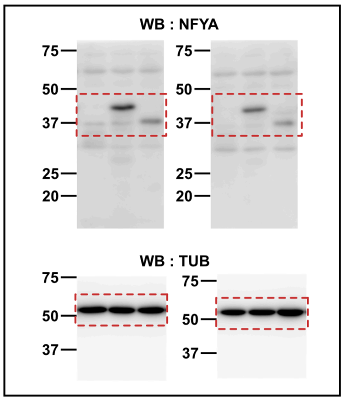

Supplementary Figure 2f

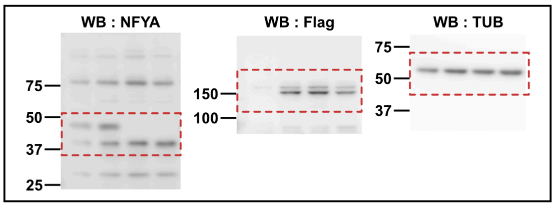

Supplementary Figure 2i

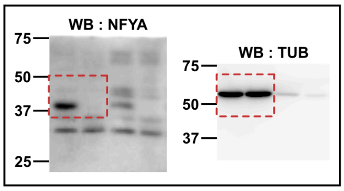

Figure 4d

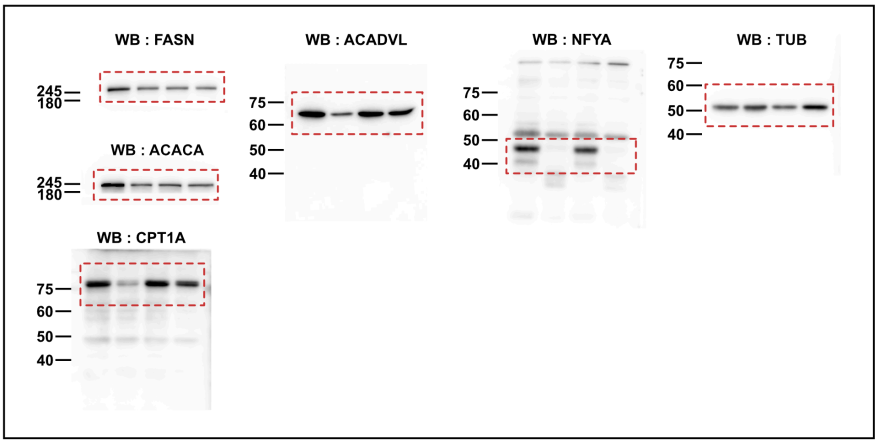

Figure 4f

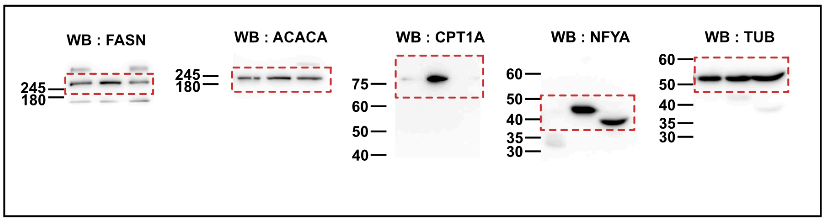

Supplementary Figure 4c

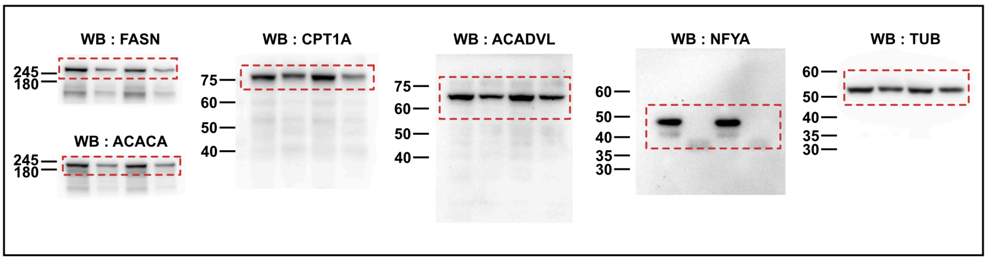

Supplementary Figure 4d

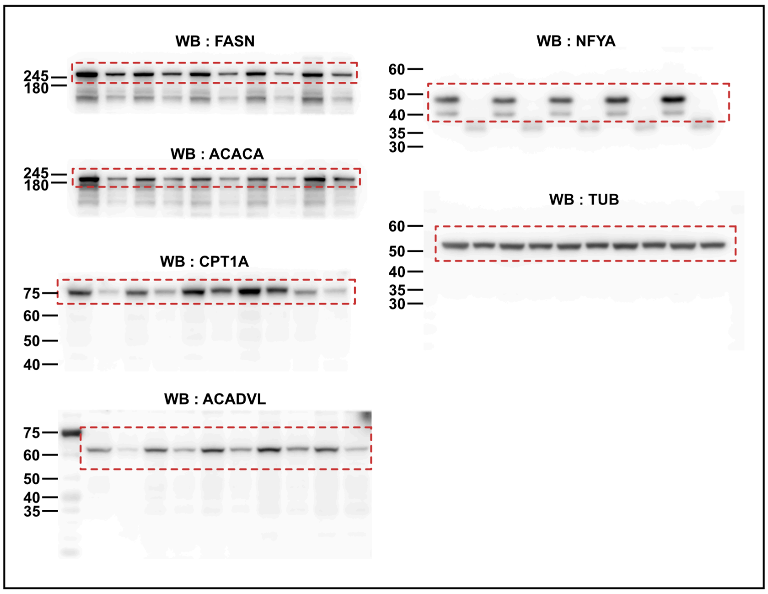

Supplementary Figure 4e

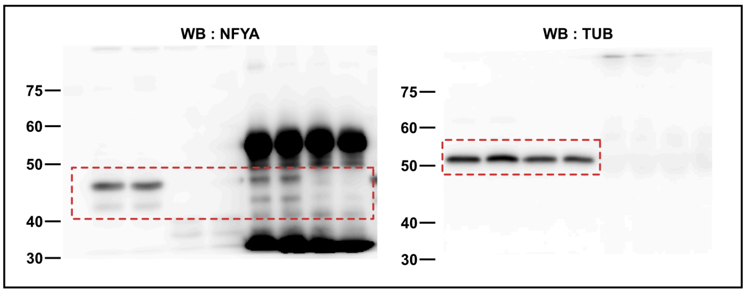

Supplementary Figure 4f

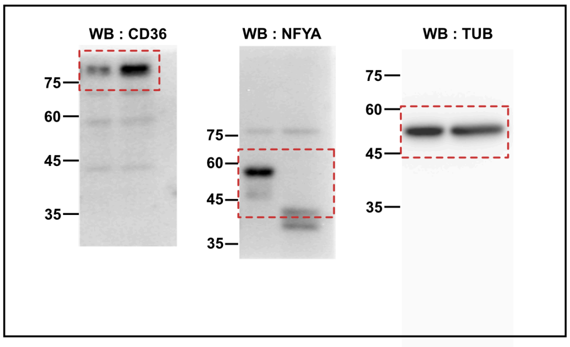

Figure 5c

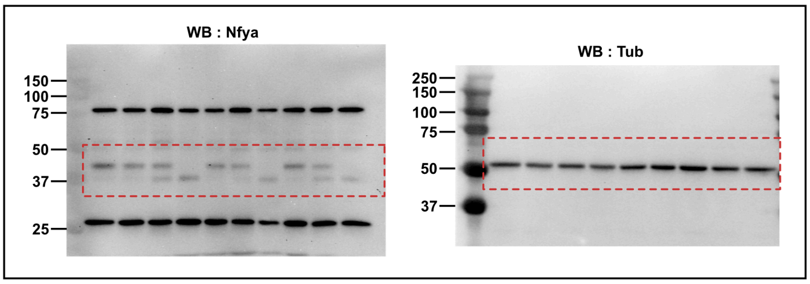

Figure 5i

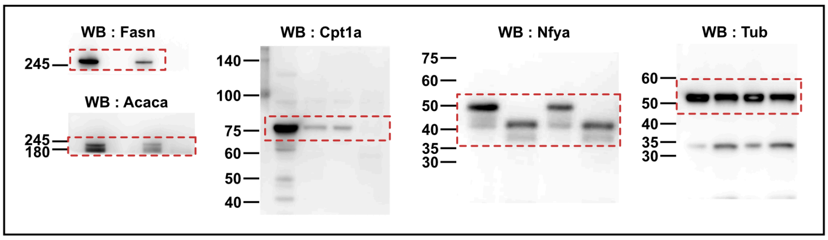

Supplementary Figure 5e

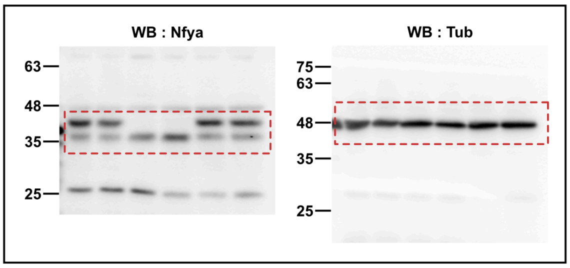

**Supplementary Fig. 6** Uncropped western blot images
